# A nanobody-based toolset to investigate the role of protein localization and dispersal in *Drosophila*

**DOI:** 10.1101/106930

**Authors:** Stefan Harmansa, Ilaria Alborelli, Emmanuel Caussinus, Markus Affolter

**Affiliations:** Growth & Development, Biozentrum, Klingelbergstrasse 50/70, University of Basel, 4056 Basel, Switzerland; Institute of Molecular Life Sciences (IMLS), University of Zurich, 8057 Zurich, Switzerland

## Abstract

Investigating the role of protein localization is crucial to understand protein function in cells or tissues. However, in many cases the role of different subcellular fractions of given proteins along the apical-basal axis of polarized cells has not been investigated *in vivo*, partially due to lack of suitable tools. Here, we present the GrabFP system, a nanobody-based toolbox to modify the localization and the dispersal of GFP-tagged proteins along the apical-basal axis of polarized cells. We show that the GrabFP system is an effective and easy-to-implement tool to mislocalize cytosolic and transmembrane GFP-tagged proteins and thereby functionally investigate protein localization along the apical-basal axis. We use the GrabFP system as a tool to study the extracellular dispersal of the Decapentaplegic (Dpp) protein and show that the Dpp gradient forming in the lateral plane of the *Drosophila* wing disc epithelium is essential for patterning of the wing imaginal disc.

## Introduction

Despite being an important property, the role of protein localization and the effects of forced protein mislocalization have not been studied extensively and hence remain in many cases not well understood. Over the last few years, genetically encoded protein binders have been introduced to basic biological research and provide novel means for protein manipulation *in vivo*. While protein function was largely studied by genetic manipulation at the DNA or RNA levels in the past, protein binders allow direct, specific and acute modification and interference of protein function *in vivo* (Kaiser, Maier et al. 2014, Bieli, Alborelli et al. 2016) and might therefore represent valid tools to study protein localisation.

Several types of protein binders exist (for recent reviews see Helma, Cardoso et al. 2015, Pluckthun 2015). One class of widely applied protein binders are the so-called nanobodies, which are derived from single chain antibodies found in members of the Camelid family. A nanobody specifically recognizing GFP (vhhGFP4, see Saerens, Pellis et al. 2005) has been extensively used for cell and developmental biology applications. Importantly, vhhGFP4 can be fused to other proteins without losing its activity and specificity *in vivo* (Rothbauer, Zolghadr et al. 2008). As a consequence, vhhGFP4 has been functionalized by fusing it to different protein domains in order to visualize (Rothbauer, Zolghadr et al. 2006), relocalize (Berry, Olafsson et al. 2016) and degrade (Caussinus, Kanca et al. 2012, Shin, Park et al. 2015) GFP-tagged proteins of interest. More recently, GFP nanobodies were used to generate inducible tools that allow controlled transcription (Tang, Szikra et al. 2013) and enzyme activity (Tang, Rudolph et al. 2015), and to generate synthetic receptors (Harmansa, Hamaratoglu et al. 2015, Morsut, Roybal et al. 2016), to mention only a few examples.

Recently, we utilized vhhGFP4 to create a synthetic receptor for GFP-tagged signalling molecules and termed it morphotrap (Harmansa, Hamaratoglu et al. 2015). Morphotrap consists of a fusion protein between vhhGFP4 and the mouse CD8 transmembrane protein, designed such that the nanobody is presented extracellularly along the surface of cells. In combination with a GFP-tagged version of the Decapentaplegic (eGFP-Dpp) morphogen, a secreted signalling molecule, morphotrap provided a powerful tool to modify secretion and extracellular dispersal of eGFP-Dpp in the *Drosophila* wing disc tissue (Harmansa, Hamaratoglu et al. 2015).

Here we introduce the GrabFP (*grab Green Fluorescent Protein*) toolbox, consisting of morphotrap and five novel synthetic GFP-traps that either localize to both the apical and basolateral compartment (morphotrap) or preferentially to one compartment: apical (GrabFP-A) or basolateral (GrabFP-B, Figure 1A). For each of these three localizations, two constructs were created in which the vhhGFP4 domain either faces the extracellular space (GrabFP_Ext_) or the intracellular milieu (GrabFP_Int_). As a consequence, the GrabFP system can interfere with target proteins in the extracellular and the intracellular space (Figure 1A).

**Figure 1.**
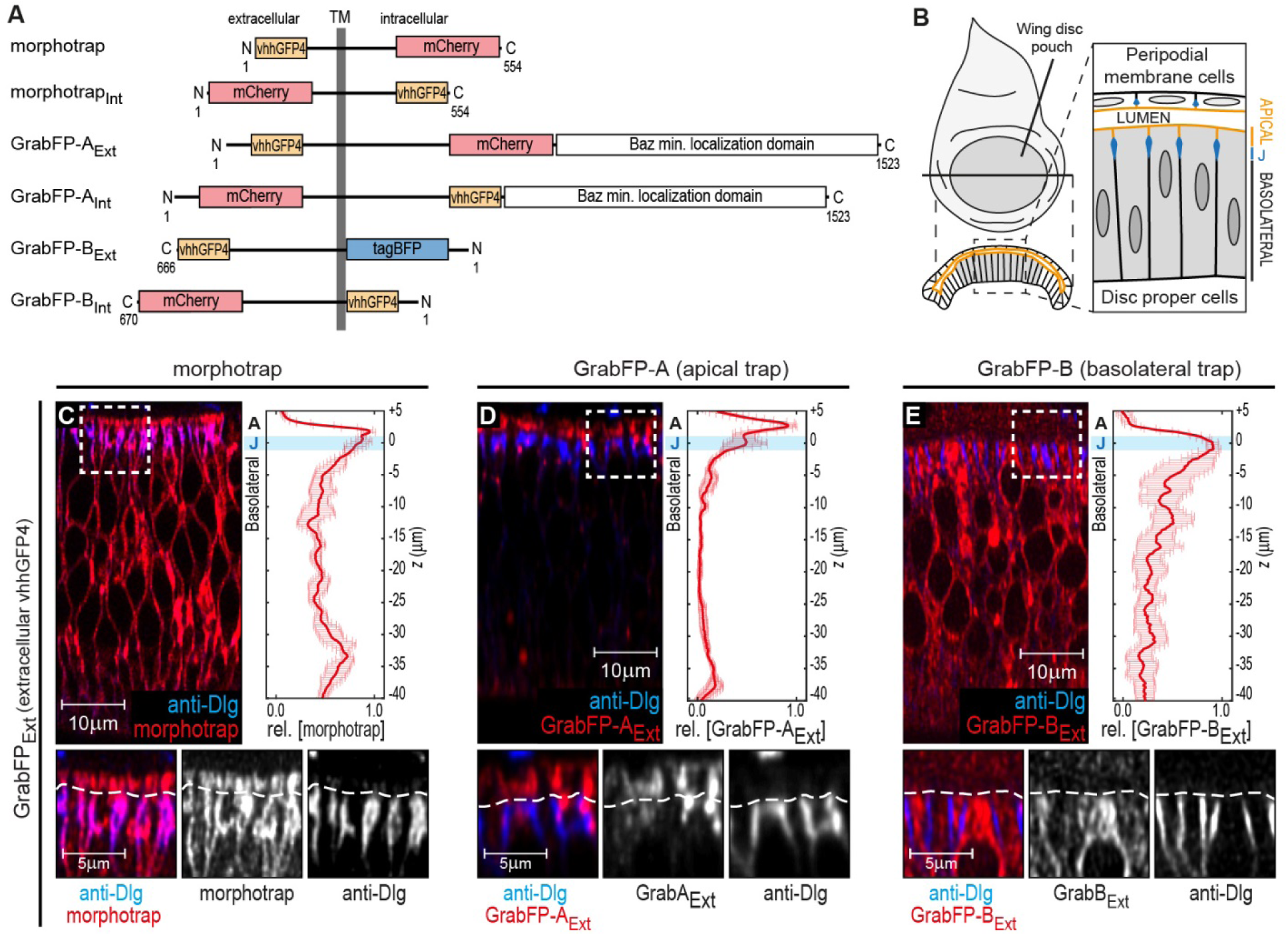
The GrabFP constructs localize to distinct regions along the apical-basal axis. **A**, Linear representation of the six different versions of the GrabFP system; the constructs exist in two topologies with the GFP-nanobody (vhhGFP4) either facing extracellular (Ext) or intracellular (Int). Numbers refer to the amino acid positions from the N-terminus (N) to the C-terminus (C). TM = transmembrane domain. **B**, Schematic representation of wing disc morphology, the junctions (J) are marked in blue. **C-E**, Cross-sections of wing discs expressing morphotrap (C), GrabFP-A_Ext_ (D) and GrabFP-B_Ext_ (E) in the wing pouch (*nub::Gal4*). The GrabFP tools are shown in red and the junctions are visualized by staining for Dlg (blue). In the magnifications the junctional level is marked by a dotted line. Relative distribution of the GrabFP tools along the A-B axis in respect to the junctions (marked by Dlg) is quantified in the plots to the right (*n*=4 for each plot, error bars represent the standard deviation). For details on the quantification see methods and Figure 1-Figure Supplement 3.

In the following, we first investigate the potential of these anchored GFP-traps to modulate protein localization along the apical-basal (A-B) axis and in a second part to modify the extracellular dispersal of specific pools of secreted signalling molecules, e.g. morphogens. Our results show that the functional Decapentaplegic (Dpp) morphogen gradient forms in the lateral plane of the wing disc epithelium.

## Results

### The GrabFP system consists of localized GFP-traps

Analogous to morphotrap, the novel GFP-traps GrabFP-A and GrabFP-B are fusion proteins consisting of vhhGFP4 fused to transmembrane proteins determining the localization and to a fluorescent protein as a marker (Figure 1A). All constructs of the GrabFP system were implemented as Gal4 and LexA-inducible transgenes (see methods).

To test the localization and function of the GrabFP system, we made use of the *Drosophila* wing imaginal disc epithelium, a well characterized model system to study epithelial polarity (Tepass 2012, Flores-Benitez and Knust 2016) and dispersal of extracellular signalling proteins, e.g. morphogens (Therond 2012, Gradilla and Guerrero 2013, Akiyama and Gibson 2015, Langton, Kakugawa et al. 2016). The wing imaginal disc consists of two contiguous, monolayered epithelial sheets, the pseudo stratified disc proper (DP) epithelium and the squamous peripodial membrane (PPM; see Figure 1B). The apical surface of both, the DP and the PPM, is facing a luminal cavity formed between them. In this study, we characterized the expression and activity of the GrabFP toolset focusing on the columnar cells of the DP epithelium, which will form the adult wing. Visualization of the junctions via the localization of the septate junction component Discs-large (Dlg, see methods) was used to mark the border separating the apical and basolateral compartment in DP cells.

In order to restrict the GFP-traps to specific regions along the A-B axis, the GFP-nanobody was fused to a fluorophore (as a visualization marker) and a scaffold protein of known subcellular localization. Morphotrap, based on the mouse CD8 protein scaffold, was shown to localize to both the apical and the basolateral domain (see Figure 1C and Harmansa, Hamaratoglu et al. 2015). The morphotrap_Int_ construct, in which the nanobody faces the cytosol, also localizes to the apical and basolateral compartments (Figure 1-Figure Supplement 1A).

In order to generate an apically anchored trap (GrabFP-A), we made use of the transcript 48 (T48) protein (Kölsch, Seher et al. 2007). However, since a fusion protein between the GFP-nanobody, T48 and mCherry showed only mild apical enrichment (not shown), we attached the minimal localization domain of Bazooka (Krahn, Klopfenstein et al. 2010) to the C-terminus of the fusion protein (see Figure 1A and methods for details). Expression in DP cells of both versions of GrabFP-A, GrabFP-A_Ext_ and GrabFP-A_Int_, resulted in strong enrichment in the apical compartment, while only low amounts of GrabFP-A_Ext_ or GrabFP-A_Int_ were observed along the basolateral domain (Figure 1D and Figure 1-Figure Supplement 1B).

GrabFP-B, a basolaterally anchored GFP-trap, is based on the Nrv1 protein (Figure 1A). Nrv1 is a subunit of the Na^+^/K^+^ ATPase (Sun and Salvaterra 1995, Xu, Sun et al. 1999), and localizes to the basolateral compartment of the wing disc even when overexpressed (Genova and Fehon 2003, Paul, Palladino et al. 2007). In DP cells, GrabFP-B_Ext_ and GrabFP-B_Int_ exclusively localized to the basolateral compartment with no detectable signal along the apical compartment (Figure 1E and Figure 1-Figure Supplement 1C).

Expression of the GrabFP constructs in the wing imaginal disc yielded viable and fertile adults with normal wings (Figure 1-Figure Supplement 2), suggesting that the GrabFP system is inert in the absence of GFP and can be used as a tool to study protein function along the A-B axis in the wing imaginal disc.

### Mislocalizing transmembrane and cytosolic proteins along the A-B axis using the GrabFP system

We wanted to test whether the interaction between our localized GFP-traps and a GFP-tagged target protein, transmembrane or cytosolic, can result in defined mislocalization of the target protein. Therefore, single components of the GrabFP system were co-expressed with different target proteins in defined domains of the wing imaginal disc (*hh::Gal4* for GrabFP_Ext_ and *ptc::Gal4* for GrabFP_Int_), while neighbouring areas were used as an internal control for the analysis of wild-type target protein localization. We analysed and measured the changes in distribution along the A-B axis of a total of 14 GFP/YFP tagged proteins, of which 10 were transmembrane and 4 were cytoplasmic proteins. We choose to use target proteins localizing either exclusively to a subcellular compartment (apical or basolateral) or, alternatively, throughout the A-B axis.

We tested the GrabFP_Ext_ system, which displays the anti-GFP nanobody along the extracellular side (Figure 2A), in combination with 8 transmembrane proteins extracellularly tagged with GFP/YFP. Expression of either GrabFP-A_Ext_ (Figure 2G) or GrabFP-B_Ext_ (Figure 2H) caused significant changes in the distribution of all 8 proteins tested, such that all target proteins acquired a novel biased distribution along the A-B axis. Generally, GrabFP-A_Ext_ efficiently induced mislocalization of target proteins (i.e. the gain of a novel apical fraction in proteins excluded from the apical compartment, as seen for NrxIV-YFP, Figure 2B) and stabilization of an existing apical fraction (as seen for Dlp-YFP, Dally-YFP, PMCA-YFP, Figure 2C and Figure 2-Figure Supplement 1A-B). However, GrabFP-A_Ext_ expression did not result in efficient depletion of the basolateral protein fraction (Figure 2G). This might be due to the fact that GrabFP-A_Ext_ itself was partially mislocalized due to interaction with polarized target proteins and showed enhanced localization to the basolateral compartment (Figure 2-Figure Supplement 1E). In contrast, GrabFP-B_Ext_ displayed a strong potential in depleting apical target-protein fractions (Figure 2D-F and Figure 2-Figure Supplement 1C-D). In particular, GrabFP-B_Ext_ significantly reduced the apical pool and increased the basolateral fraction of Dally-YFP, Notch-YFP, Fra-YFP, Crb-GFP and Ed-YFP (Figure 2H). Furthermore, GrabFP-B_Ext_ was resistant to mislocalization induced by target protein-interaction (Figure 2-Figure Supplement 1F).

**Figure 2.**
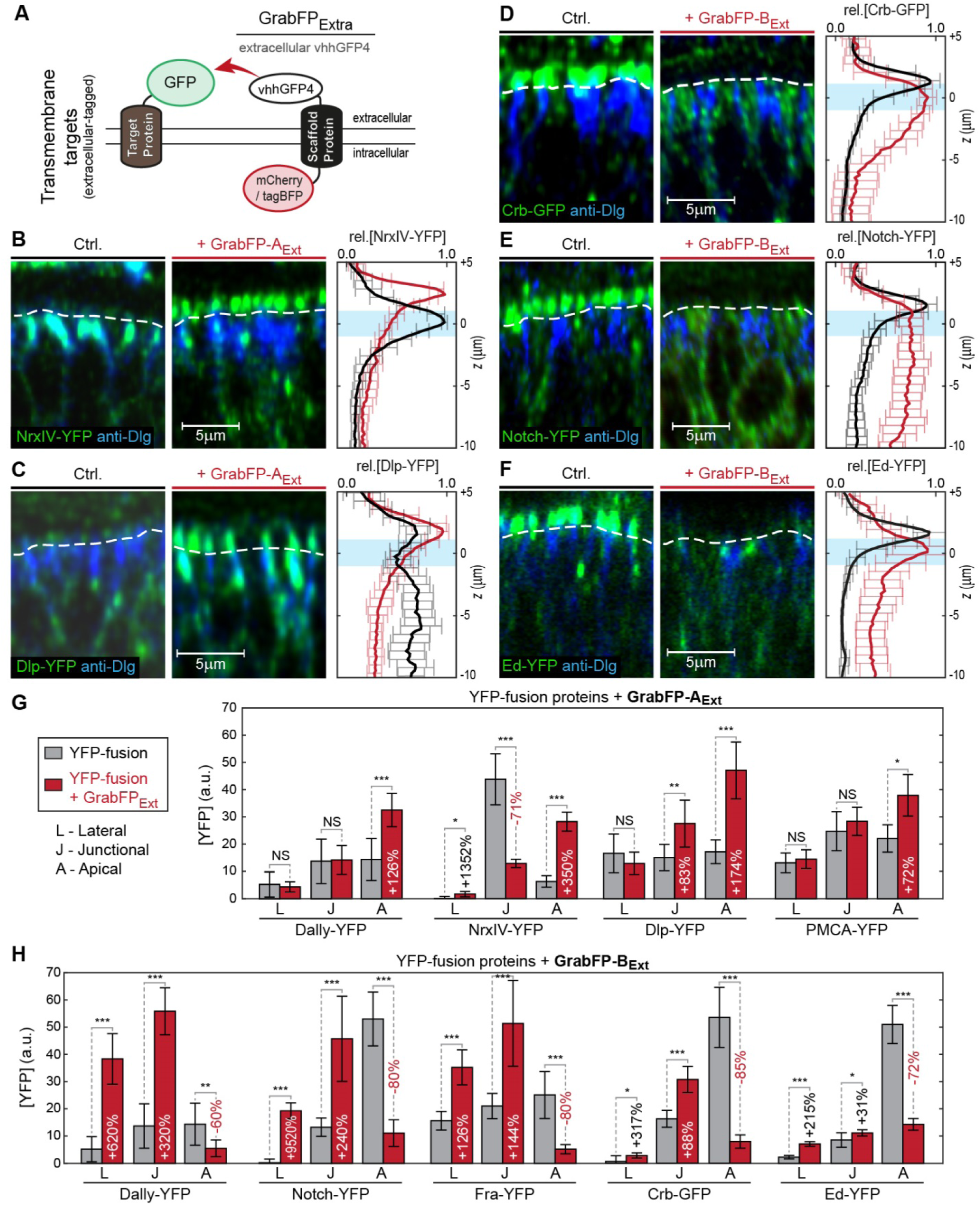
Mislocalization of transmembrane proteins using the GrabFPExt system. **A**, In the GrabFP_Ext_ system the GFP-nanobody (vhhGFP4) faces the extracellular space and can interact with extracellular-tagged transmembrane proteins. **B-C**, Optical cross-section of wing disc cells expressing either NrxIV-YFP (B) or Dlp-YFP (C) alone (Ctrl., left) or together with GrabFP-A_Ext_ (middle). The junctional level is marked by a dotted line. Quantification of relative target-protein localization (right) along the A-B axis in the absence (black) or in the presence of GrabFP-A_Ext_ (red). The position of the junctions is marked by a blue bar. (Error bars show the standard deviation). **D-F**, Optical cross sections showing the localization of Crb-GFP (D), Notch-YFP (E) or Ed-YFP (F) in the absence (left) or in the presence of GrabFP-B_Ext_ (middle). Quantifications are shown to the right. **G**, Quantification of target-protein concentrations in the apical (A), junctional (J) and basolateral (L) compartments in control conditions (grey) or when co-expressed with GrabFP-A_Ext_ for all tested GrabFP target-protein interactions. For representative optical cross-sections of the not shown target-proteins see Figure 2-Figure Supplement 1A-B. NS = non-significant. **H**, Quantification of protein concentration changes along the A-B axis induced by GrabFP-B_Ext_ co-expression (also see Figure 2-Figure Supplement 1C-D). (Sample numbers for plots in B-F and quantifications in G-H: Dally *n*=10, NrxIV *n*=10, Dlp *n*≥8, PMCA *n*=5, Notch *n*≥8, Fra *n*≥8, Crb *n*=8, Ed *n*≥6, significance was assessed using a two-sided Student’s *t*-test with unequal variance, * *p*<0.05, ** *p*<0.005, *** *p*<0.0005)

In summary, expression of GrabFP_Ext_ components leads to significant mislocalization of target proteins with an average apical enrichment of 2.8-folds using GrabFP-A_Ext_ and a 12.6-fold average basolateral enrichment using GrabFP-B_Ext_ (Figure 2G-H). Moreover GrabFP-B_Ext_ caused significant and efficient depletion of the apical fractions of all proteins analysed.

In a next step, we tested the mislocalization potential of the GrabFP_Int_ system, in which the anti-GFP nanobody localizes intracellularly (Figure 3A). To this aim, we used 3 transmembrane proteins (Fat, Nrv1, Nrv2) containing an intracellular GFP/YFP tag and 3 GFP/YFP-tagged cytoplasmic proteins (Arm, αCat, Hts). We observed significant changes in the distribution of both transmembrane and cytoplasmic target proteins (Figure 3G-H). GrabFP-B_Int_ efficiently depleted the apical fraction of Fat-GFP and induced strong enrichment of its basolateral fraction (Figure 3B). In contrast, GrabFP-B_Int_ was less efficient in mislocalizing and depleting the apical fraction of the cytoplasmic proteins αCat-GFP and Arm-GFP (Figure 3C and Figure 3-Figure Supplement 1). Concomitantly, GrabFP-B_Int_ showed a higher tendency to be mislocalized when co-expressed with these two cytosolic targets (Figure 3-Figure Supplement 1C). GrabFP-B_Int_ caused 5.7-folds of basolateral target enrichment on average. GrabFP-A_Int_ efficiently mislocalized target proteins by decreasing their basolateral concentration and increasing their apical fraction of 14-folds on average (Figure 3H). Notably, all proteins tested in combination with GrabFP-A_Int_ had a strong bias towards the basolateral side in wild-type conditions and acquired a strong apical fraction when co-expressed with GrabFP-A_Int_ (Figure 3D-F). Furthermore, GrabFP-A_Int_ showed to be resistant to mislocalization induced by target protein interaction (Figure 3-Figure Supplement 1B).

**Figure 3.**
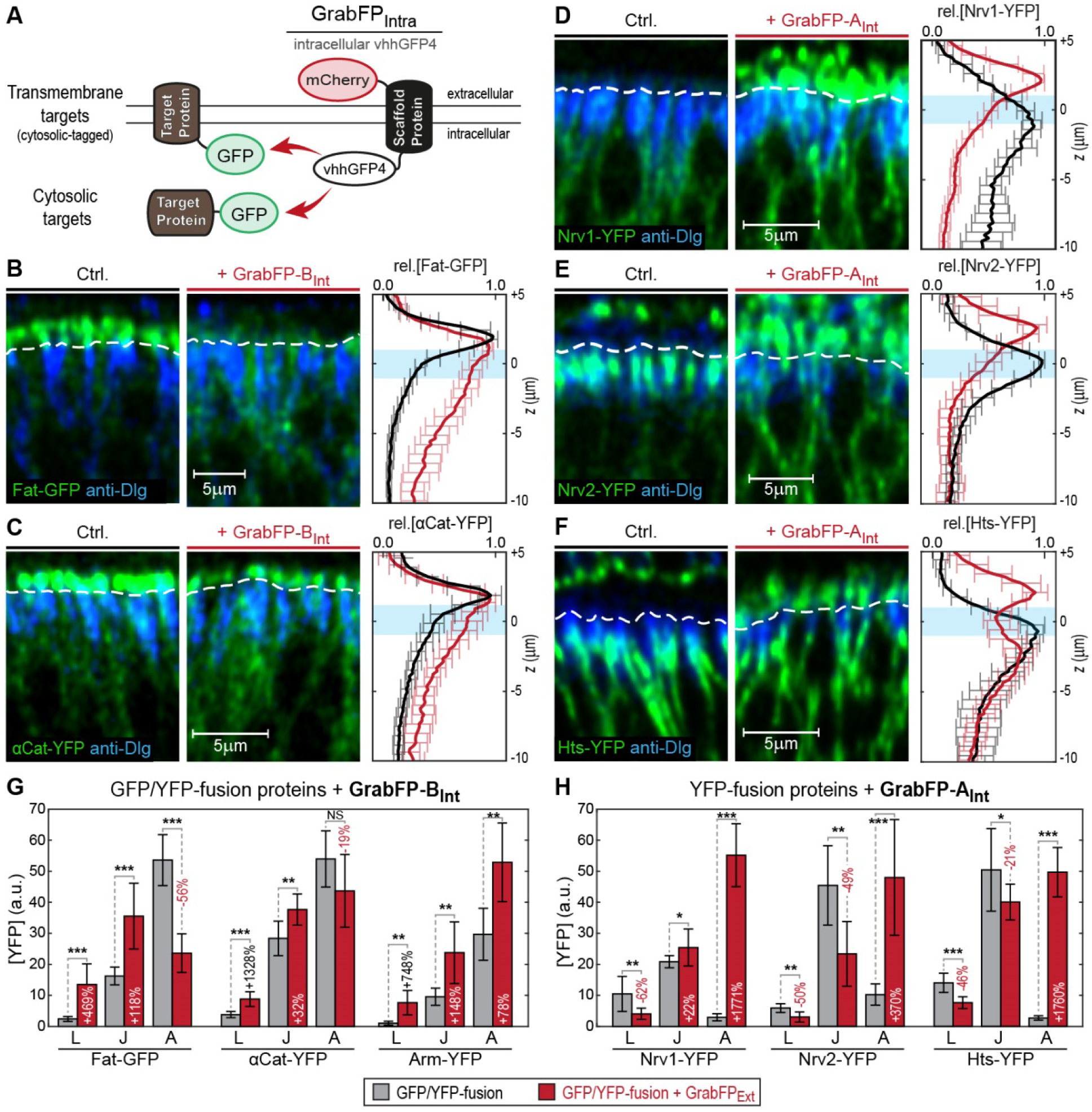
Mislocalization of GFP/YFP-tagged proteins using the GrabFP_Int_ system. **A**, With the GFP-nanobody facing the cytosol, the GrabFP_Int_ system can interact with cytosolic proteins and transmembrane proteins tagged along their cytosolic portion. **B-C**, Optical cross-sections of wing disc cells expressing either Fat-GFP (B) or αCat-YFP (C) alone (Ctrl., left) or together with GrabFP-B_Int_ (middle). A dotted line marks the junctional level. Quantification of relative target-protein localization (right) along the A-B axis in the absence (black) or in the presence of GrabFP-B_Int_ (red). The position of the junctions is marked by a blue bar. (Error bars show the standard deviation). **D-F**, Optical cross-sections showing the localization of Nrv1-YFP (D), Nrv2-YFP (E) or Hts-YFP (F) in the absence (left) or in the presence of GrabFP-A_Int_ (middle). Quantifications are shown to the right. **G**, Target-protein fluorescence intensities in the apical (A), junctional (J) and basolateral (L) compartments in control conditions (grey) or when co-expressed with GrabFP-B_Int_. For a representative optical cross-sections Arm-YFP see Figure 3-Figure Supplement 1A. NS = non-significant. **H**, Quantification of protein fluorescent intensity changes along the A-B axis induced by GrabFP-A_Int_ co-expression. (Sample numbers for plots in B-F and quantifications in G-H: Fat *n*=10, αCat *n*=9, Arm *n*=8, Nrv1 *n*=10, Nrv2 *n*=10, HTS *n*=10, significance was assessed using a two-sided Student’s *t*-test with unequal variance, * *p*<0.05, ** *p*<0.005, *** *p*<0.0005)

To further validate the GrabFP system as a tool to study the role of protein localization *in vivo*, we attempted to mislocalize *spaghetti squash (sqh)*, the *Drosophila* regulatory light chain of Myosin II. We made use of a Sqh-GFP transgene expressed under the control of the *sqh* promoter (*sqhSqh-GFP* flies, Royou, Field et al. 2004) that rescues the *sqh^A×4^* null allele. *Drosophila* Sqh is crucial for morphogenesis and control of epithelial cell shape (Young, Richman et al. 1993, Kiehart, Galbraith et al. 2000). Sqh-GFP is a cytosolic protein that localizes to the subapical cortex in wing disc cells (Figure 4A) and is required for maintaining the elongated shape of DP cells (Widmann and Dahmann 2009). To test whether mislocalization of Sqh-GFP from the apical cortex to the basolateral domain indeed affects DP cell shape, we expressed GrabFP-B_Int_ in *sqhSqh-GFP* flies. Expression of GrabFP-B_Int_ in *sqhSqh-GFP* female flies that are heterozygous for *sqh^A×4^* (and hence, carry one wild-type and one GFP-tagged copy of Sqh) resulted in increased Sqh-GFP levels in the basolateral domain and concomitant reduction in the basal cell surface (Figure 4B-C). In *sqhSqh-GFP* male flies, which are hemizygous for *sqh^A×4^* (and in which Sqh-GFP represents the only source of Sqh protein), Sqh-GFP mislocalization with GrabFP-B_Int_ caused an even more drastic alteration of cell shape (Figure 4D) visible as a strong constriction of the basolateral domain accompanied by a significant expansion of the apical cell surface (Figure 4F-G). This behaviour could be explained by loss of apical tension (due to the reduction of apical Sqh-GFP) and increased basolateral tension (due to mislocalized Sqh-GFP) (Figure 4E). In conclusion, GrabFP-B_Int_ altered the localization of Sqh-GFP, presumably causing significant alterations in the force distribution along the cortex of DP cells.

**Figure 4.**
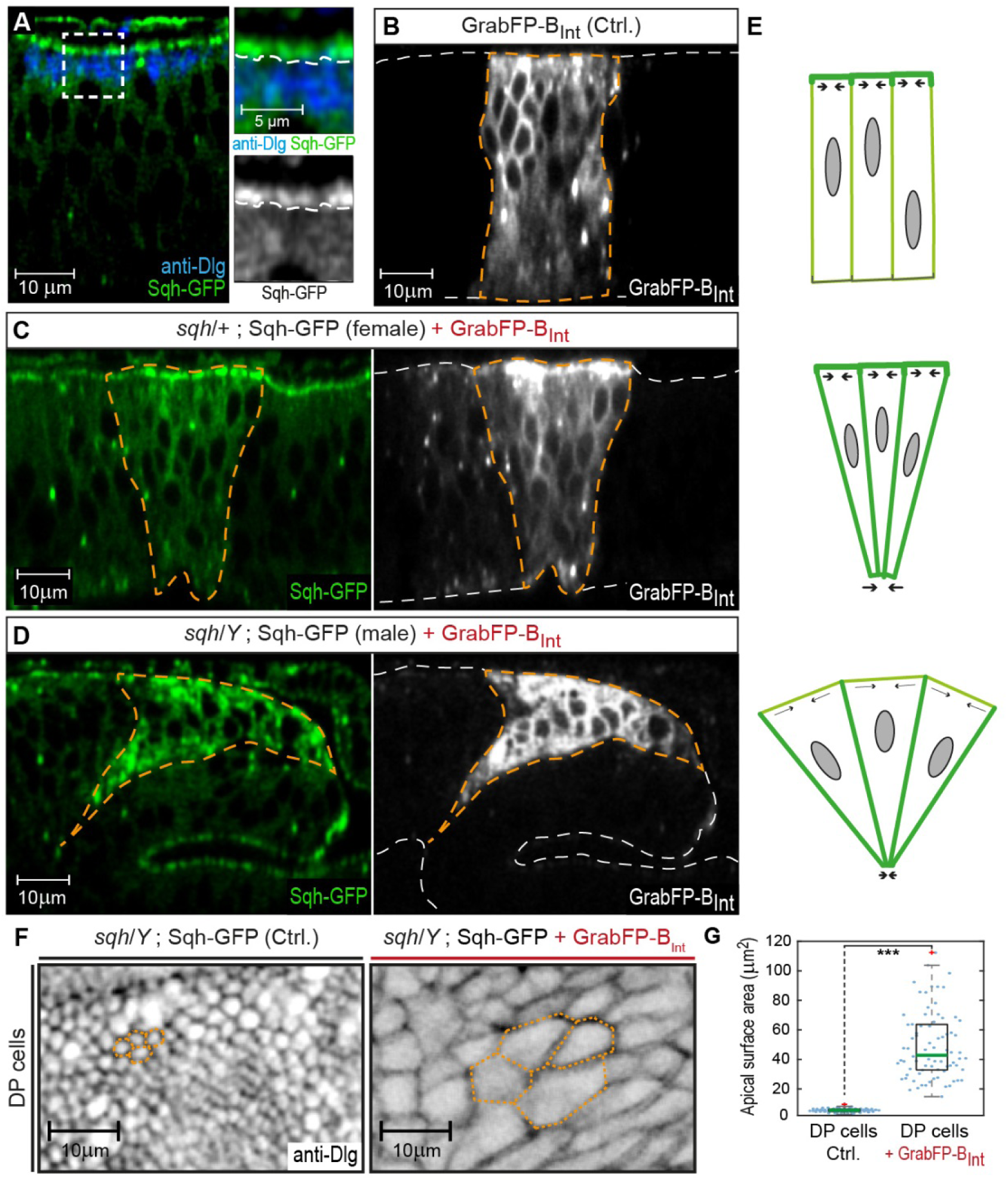
GrabFP_Int_ mediated Sqh-GFP mislocalization results in changes of DP cell shape. **A**, Optical cross-section of a wing disc expressing Sqh-GFP (green), stained for Dlg (blue). In the magnifications the junctional level is marked by a dotted line. **B-D**, Optical cross-sections of wing discs expressing GrabFP-B_Int_ (grey) in the patched domain (marked by dotted orange line, *ptc::Gal4*) either alone (Ctrl., B) or together with Sqh-GFP (green) in heterozygous *sqh* females (C) and hemizygous *sqh* males (D). Sqh-GFP mislocalization causes a drastic increase of basolateral Sqh-GFP (C-D). Mislocalization of Sqh-GFP causes cell shape alterations resulting in a triangular shape of the *ptc* domain (C-D), compared to the rectangular shape of the *ptc* domain in control discs (B). The white dotted line marks the apical (top) and basal (bottom) surface of DP cells. **E**, Schematic representation of the effect of Sqh-GFP mislocalization. Tension is higher in the apical cortex of columnar cells due to polarization of myosin II activity (top). Mislocalization of Sqh-GFP causes increased basolateral tension, leading to constriction of the basolateral cell area (middle). In *sqh* hemizygous conditions the apical surface expands, due to decreased apical myosin II activity (bottom). **F**, Projections of the junctional level of the DP columnar epithelium of the genotype shown in (D) either in the absence of GrabFP-B_Int_ (left, normal Sqh::GFP localization) or in the presence of GrabFP-B_Int_ (right, mislocalized Sqh::GFP). **G**, Quantification of apical surface area as marked in (F). The green line marks the median, statistical significance was assessed using a two-sided Students t-test (*** *p*<0.0005).

In summary, our results show that the GrabFP system offers a novel toolbox to modify protein localization along the A-B axis in a controlled manner and to study the role of protein localization and forced protein mislocalization *in vivo*.

### GrabFP as a tool to study the dispersal of the Decapentaplegic morphogen

Another potential application of the GrabFP system is to study how morphogen gradients form and control patterning and growth during animal development. Morphotrap has previously been used to address the requirement of the Dpp morphogen gradient for patterning and growth of the wing imaginal disc (Harmansa, Hamaratoglu et al. 2015). We wanted to extend these studies using the newly generated tools reported here.

A key property that has not been studied in detail is the dispersal of functional Dpp in the wing disc tissue with regard to the A-B axis. We therefore utilize the GrabFP_Ext_ system in combination with an eGFP-tagged version of Dpp (eGFP-Dpp, Teleman and Cohen 2000) to study the localization of the functional Dpp gradient along the A-B axis.

### Dpp disperses in the apical and in the basolateral compartment

In the developing wing imaginal disc, Dpp is expressed and secreted from a central stripe of anterior cells adjacent to the anterior/posterior (A/P) compartment boundary from where it forms a concentration gradient into the surrounding target tissue. The Dpp gradient in the wing disc has been visualized using different GFP-Dpp fusion proteins (Entchev, Schwabedissen et al. 2000, Teleman and Cohen 2000) and by antibody staining against endogenous Dpp protein (Gibson, Lehman et al. 2002, Akiyama and Gibson 2015). Dpp was observed in the lateral plane of the wing disc epithelial cells (Teleman and Cohen 2000) as well as apically in the wing disc lumen (Entchev, Schwabedissen et al. 2000, Gibson, Lehman et al. 2002). However, the results of these different studies were not entirely consistent and hence the routes of Dpp dispersal remain controversial.

To investigate the localization of Dpp in the wing disc, we used an eGFP-Dpp fusion protein that was shown to rescue the *dpp* mutant phenotype (Harmansa, Hamaratoglu et al. 2015). When eGFP-Dpp is expressed in its endogenous expression domain using the LexA/LOP binary expression system (*dpp::LG*, Yagi, Mayer et al. 2010), it forms a wide concentration gradient into the target tissue (Figure 5A and Harmansa, Hamaratoglu et al. 2015). In order to better characterize the localization of eGFP-Dpp in the wing imaginal disc, we acquired high-resolution confocal stacks along the z-axis. Optical cross-sections revealed that eGFP-Dpp localized prominently to dotted structures along the lateral region of the DP (Figure 5B, arrowheads), which were suggested to represent endocytic vesicles (Teleman and Cohen 2000). We did not detect eGFP-Dpp signal within the luminal space (Figure 5B, magnification). These results suggest that, using fluorescence microscopy, Dpp is prominently detected within the lateral plane of the DP epithelium.

**Figure 5.**
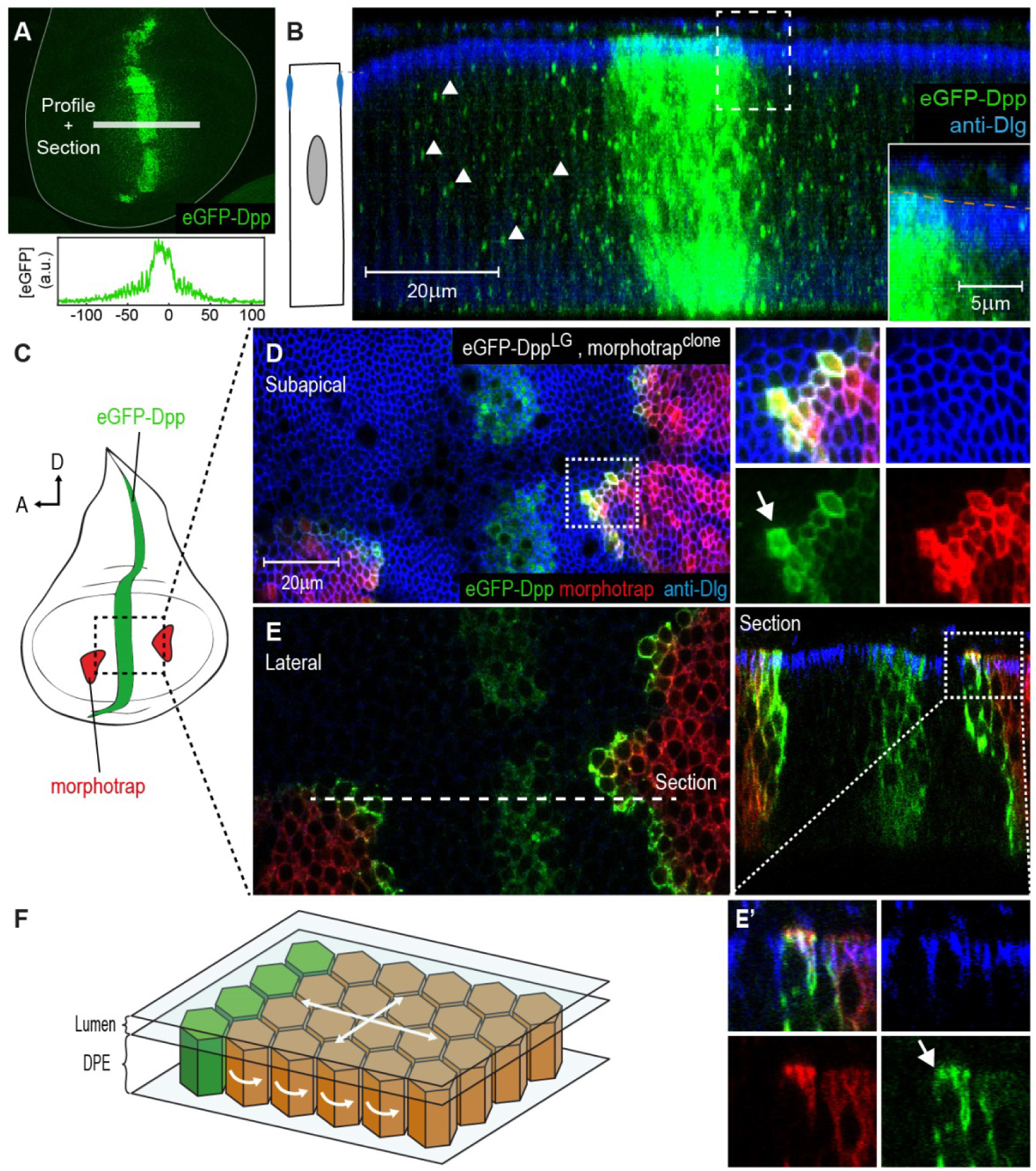
The Dpp morphogen spreads in the apical and basolateral compartment. **A**, Wing disc expressing eGFP-Dpp in the central Dpp stripe and eGFP fluorescence profile (bottom). **B**, Optical cross-section of a wing disc as shown in (A) additionally stained for Dlg (blue). eGFP-Dpp is prominently observed in spots (arrowheads) along the lateral axis of the disc but not in the wing disc lumen (see magnified insert). **C**, Scheme of morphotrap expression in clones and eGFP-Dpp in the central *dpp* stripe. **D**, Subapical projection of a wing disc expressing eGFP-Dpp in the *dpp* stripe and two lateral morphotrap clones. Magnifications to the right show apical eGFP-Dpp immobilization on the proximal surface of morphotrap clones. **E**, Lateral projection of the wing disc shown in (D). An optical cross-section to the right shows low level apical (also see arrow in magnification in (E’)) and high level basolateral immobilization of eGFP-Dpp. **F**, Scheme of the wing disc epithelium. The extracellular environment in which Dpp disperses differs between the luminal cavity and the lateral plane of the epithelium (scheme adapted from (Umulis and Othmer 2013)).

Morphotrap was reported to immobilize and accumulate eGFP-Dpp on the cell surface (Harmansa, Hamaratoglu et al. 2015). Therefore, we used morphotrap to visualize even low levels of extracellular eGFP-Dpp and to determine where along the A-B axis eGFP-Dpp encounters morphotrap-expressing target cells. When we expressed eGFP-Dpp in its central stripe source (using *dpp::LG*) and morphotrap in clones (Figure 5C), we observed high amounts of immobilized eGFP-Dpp on the proximal surface (the one facing the source of Dpp) of morphotrap clones situated in the target tissue (Figure 5D-E). Subapical projections (Figure 5D) as well as optical cross sections (Figure 5E’) showed that low amounts of eGFP-Dpp accumulated on the apical surface of morphotrap clones. However, the prominent majority of eGFP-Dpp accumulation was observed along the basolateral cell surface of morphotrap clones (Figure 5E). These results suggest that low amounts of eGFP-Dpp disperse in the apical/luminal compartment while the majority of eGFP-Dpp dispersal takes place along the basolateral compartment.

### GrabFP can specifically interfere with sub-fractions of the Dpp gradient

To investigate the role of apical and basolateral Dpp pools in patterning and growth control, we expressed eGFP-Dpp in the stripe source (using *dpp::LG*) and the different versions of the GrabFP_Ext_ system in the posterior compartment (using *hh::Gal4*, see Figure 6B-D, left). Thereby we specifically interfered with Dpp dispersal in the posterior compartment, not modifying Dpp production and secretion.

**Figure 6.**
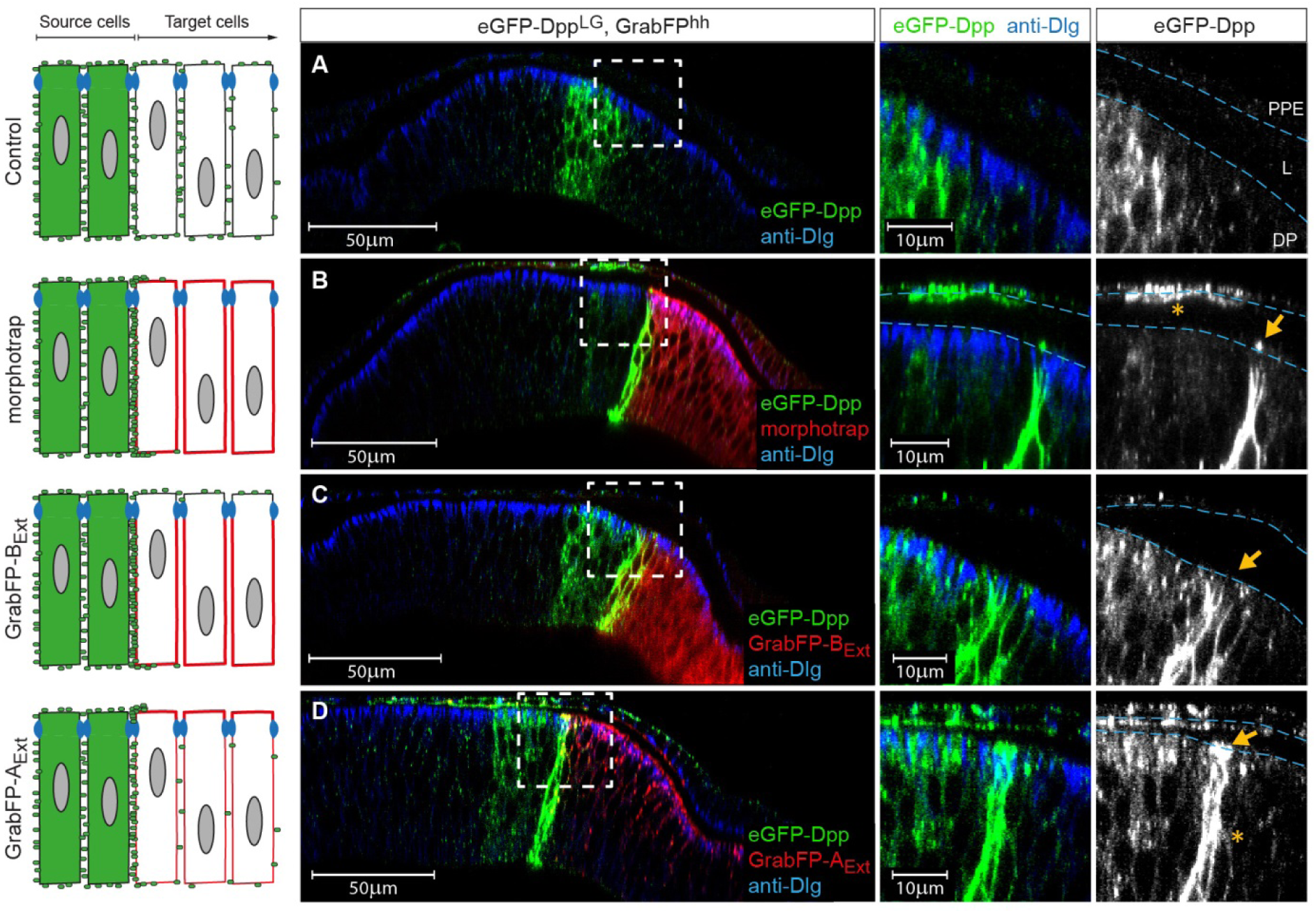
The GrabFP_Ext_ system can interfere with specific sub-fractions of the Dpp morphogen gradient. Optical cross sections of wing discs either expressing eGFP-Dpp (green) in the stripe source (A) or eGFP-Dpp in the stripe and the different versions of the GrabFP system (red, B-D) in the posterior compartment of disc proper and PPM cells (*hh::Gal4*). **A**, When expressed alone (Ctrl.), eGFP-Dpp is mainly observed in the lateral plane of the DP epithelium. Peripodial epithelium (PPE), lumen (L) and disc proper epithelium (DP). **B**, Posterior expression of morphotrap results in strong eGFP-Dpp immobilization along the basolateral domain and low or no apical immobilization (see arrow in the magnification to the right). eGFP-Dpp is also immobilized on the apical surface of PPM cells overlaying the Dpp DP source (see asterisk in magnification). **C**, Posterior expression of GrabFP-B_Ext_ results in exclusive immobilization of eGFP-Dpp in the basolateral domain. No apical immobilization is observed, neither in DP (see arrow) nor PPM cells. **D**, Expression of GrabFP-A_Ext_ in the posterior compartment results in strong basolateral (asterisk) and apical (arrow) immobilization of eGFP-Dpp.

As shown above, eGFP-Dpp expressed in a wild type background is observed in presumptive vesicular structures along the lateral plane of the epithelium, but is not present at detectable levels in the wing disc lumen (Figure 6A). Posterior morphotrap expression resulted in immobilization of eGFP-Dpp predominantly along the basolateral compartment of target cells adjacent to the Dpp source. In few cases eGFP-Dpp immobilization was observed along the apical surface of morphotrap expressing cells (see Figure 6B, arrow in right image and Figure 6-Figure Supplement 1A). Since the A/P boundary in the PPM is shifted anteriorly, morphotrap is also expressed in the PPM cells covering the Dpp DP source. Interestingly, PPM cells covering the Dpp DP source showed substantial immobilized eGFP-Dpp on their luminal surface (Figure 6B, asterisk in right image). This observation suggests that a fraction of Dpp is secreted into the lumen and disperses in the luminal cavity. These results show that posterior expression of morphotrap reduces spreading of apical and basolateral Dpp pools into the posterior compartment.

Posterior expression of GrabFP-B_Ext_ resulted in the exclusive basolateral immobilization of eGFP-Dpp close to the source (Figure 6B), consistent with its restricted localization to the basolateral membrane.

In sharp contrast, posterior expression of GrabFP-A_Ext_ resulted in strong apical and peripodial, but also basolateral immobilization of eGFP-Dpp (Figure 6D and Figure 6-Figure Supplement 1C). Therefore, it seems that the relative small portion of GrabFP-A_Ext_ localizing to the basolateral side is large enough to interfere with basolateral eGFP-Dpp dispersal (or that eGFP-Dpp relocalizes GrabFP-A_Ext_). The increased levels of apical eGFP-Dpp immobilization might also hint towards mislocalization of basolateral immobilized eGFP-Dpp to the apical compartment by GrabFP-A_Ext_.

In summary, the GrabFP_Ext_ system can be used to interfere with both apical and basolateral dispersal (morphotrap) or to specifically interfere with basolateral eGFP-Dpp dispersal (GrabFP-B_Ext_). However, localization of GrabFP-A_Ext_ is not specific enough to exclusively interfere with apical Dpp dispersal (see also Discussion).

### Basolateral Dpp dispersal is required for patterning and growth of the *Drosophila* wing

In an earlier study using morphotrap we reported that Dpp dispersal is important for wing disc growth and patterning (Harmansa, Hamaratoglu et al. 2015). Since we find that Dpp is prominently found in the basolateral compartment, we wanted to use the newly generated GrabFP system to investigate whether basolateral Dpp dispersal is required for patterning of the wing. We therefore compared the p-Mad signalling response of *dpp^d8/d12^* mutant wing discs rescued with eGFP-Dpp (normal Dpp dispersal) to *dpp^d8/d12^* mutant wing discs rescued with eGFP-Dpp expressing either morphotrap (apical and basolateral Dpp dispersal reduced) or GrabFP-B_Ext_ (only basolateral Dpp dispersal reduced) in the posterior compartment, respectively (Figure 7A-H).

**Figure 7.**
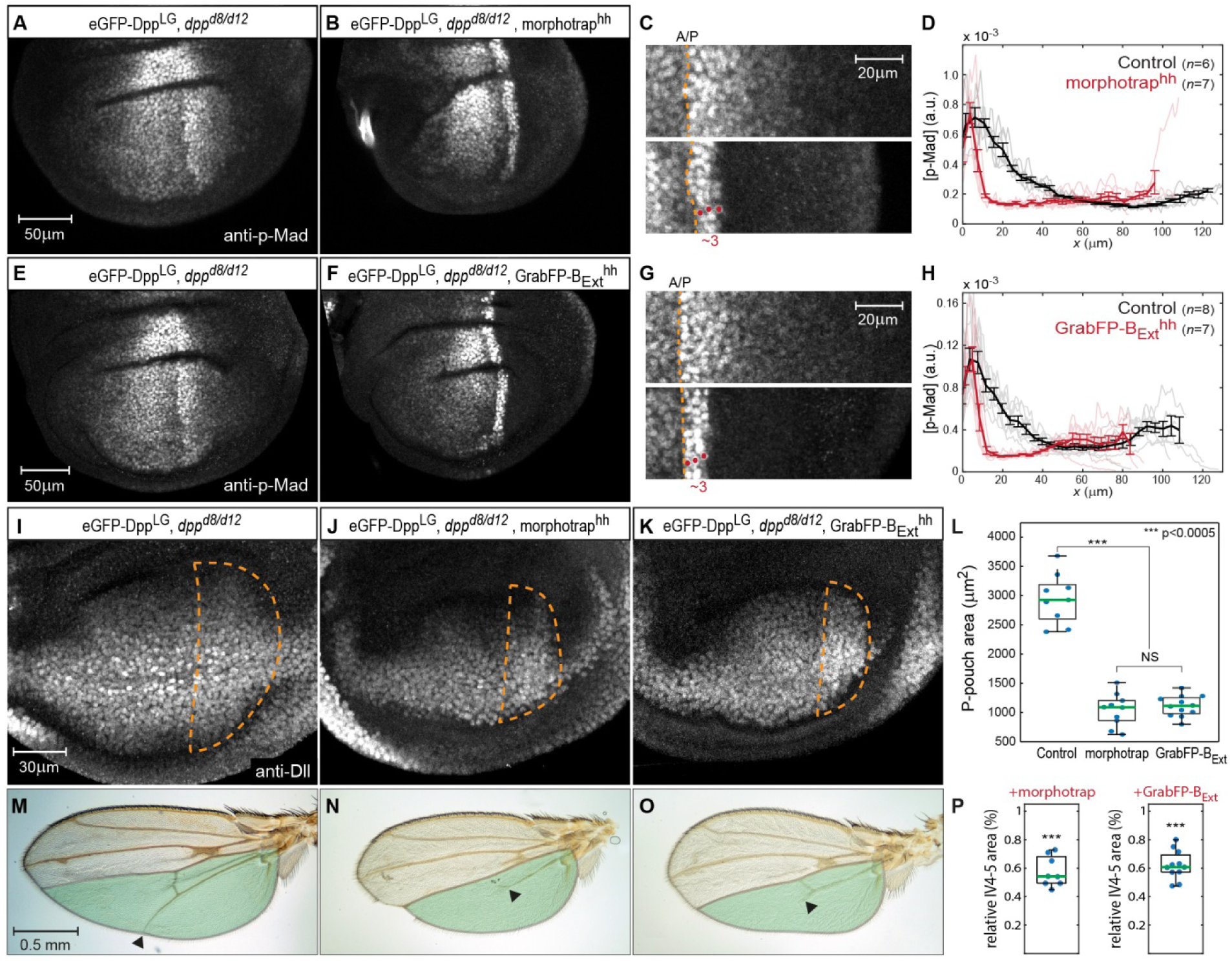
Basolateral Dpp spreading is required for patterning and size control. **A-B**, p-Mad staining in representative *dpp^d8/d12^* mutant wing disc rescued by eGFP-Dpp (A) and in *dpp^d8/d12^* wing disc rescued by eGFP-Dpp expressing morphotrap in the posterior compartment (*hh::Gal4*, B). **C**, Magnifications of the posterior, dorsal pouch region of the images shown in (A-B). The A/P boundary is marked by a dotted yellow line. **D**, Average posterior p-Mad profiles of 98-100h AEL old *dpp^d8/d12^* wing disc rescued by eGFP-Dpp (black) and *dpp^d8/d12^* wing disc rescued by eGFP-Dpp expressing morphotrap (red). **E-H**, Representative wing discs and quantification of p-Mad levels in *dpp^d8/d12^* wing disc rescued by eGFP-Dpp (E, black in H) and *dpp^d8/d12^* wing disc rescued by eGFP-Dpp expressing GrabFP-B_Ext_ in the posterior compartment (F, red in H). **I-K**, Representative 98-100h AEL old wing discs of the indicated genotypes stained for Distal-less (Dll) as a marked for pouch size. The posterior wing pouch is outlined by a dotted yellow line. **L**, Quantification of posterior wing pouch area as shown in (I-K). (Control *n*=9, morphotrap *n*=10, GrabFP-B_Ext_ *n*=12) **M-O**, Female wings of the genotypes indicated. The area posterior to vein 4 (IV4-5) is marked in green. Block of apical and basolateral, as well as block of basolateral Dpp dispersal results in a loss of the distal parts of wing vein 5 and hence patterning (see arrowheads). **P**, Quantification of relative IV4-5 area as indicated in (M-O). (*** *p*>0.0005, Control *n*=10, morphotrap *n*=8, GrabFP-B_Ext_ *n*=11)

In control conditions (normal Dpp spreading), p-Mad forms a wide bilateral concentration gradient into the anterior and posterior compartment (>40μm; see Figure 7A). In contrast, reduction of apical and basolateral spreading by expression of morphotrap in the posterior compartment resulted in a drastic reduction of the posterior p-Mad range to ~3 cells or ~10μm (Figure 7B-D). Interestingly, specifically interfering with basolateral Dpp spreading by posterior expression of GrabFP-B_Ext_ also resulted in a reduction of posterior P-Mad range to ~3 cells or ~10μm (Figure 7F-H), a result strikingly similar to the morphotrap experiment. Hence, these experiments demonstrated that basolateral Dpp spreading is required for proper Dpp signalling range and patterning and that apical/luminal Dpp spreading is not sufficient.

We also investigated whether growth of the wing disc requires basolateral Dpp spreading. Indeed, we found that the posterior wing pouch area visualized by immunostaining against Distal-less (Dll) was reduced to a similar extend when expressing either morphotrap or GrabFP-B_Ext_ in the posterior compartment (Figure 7I-L). Accordingly, the posterior wing blade area was reduced to a similar extend in both the morphotrap and the GrabFP-B_Ext_ condition (Figure 7M-P). In addition, and consistent with the strongly reduced p-Mad range, the distal portion of wing vein 5 was lost upon posterior expression of morphotrap or GrabFP-B_Ext_ (19/19 wings).

In summary, these results show that basolateral, not apical/luminal Dpp dispersal is important for patterning and size control of the wing disc and the adult wing. To further test the requirement of luminal Dpp spreading, we expressed morphotrap in PPM cells to hinder luminal Dpp dispersal (Figure 7-Figure Supplement 1). However, we observed only very mild effects on wing patterning and growth in this condition, supporting the view that apical Dpp spreading plays a minor role in wing development.

### Dpp dispersal in the basal and lateral plane control wing disc growth

Our results suggest a prominent role of basolateral Dpp spreading in the wing imaginal disc. Hence, we wanted to further dissect the function of Dpp spreading along the basolateral compartment. The basolateral compartment consists of the lateral region, where epithelial cells are compactly surrounded by their neighbours, and the basal region, where cells contact the extracellular matrix (ECM) of the basal lamina (BL). Dpp is known to interact with the heparin sulphate proteoglycans Dally and Dally-like localizing to the apical and lateral region (Figure 2C and Figure 2-Figure Supplement 1A) as well as with Collagen IV localizing to the BL (Wang, Harris et al. 2008).

In order to investigate the role of Dpp spreading in the lateral plane versus Dpp spreading in the BL, we generated GrabFP-ECM, a GFP-trap localizing to the extracellular matrix of the BL. GrabFP-ECM is a fusion protein consisting of the coding sequence of the *Drosophila* Collagen IV gene *viking* (*vkg*) (Yasothornsrikul, Davis et al. 1997, Wang, Harris et al. 2008), vhhGFP4 and mCherry inserted between the first and the second exon of *vkg* (see Methods). When expressed in the larval fat body (*r4::Gal4*), GrabFP-ECM was secreted into the haemolymph, distributed throughout the larval body and integrated into the BL of the wing disc (Figure 8A-B).

**Figure 8.**
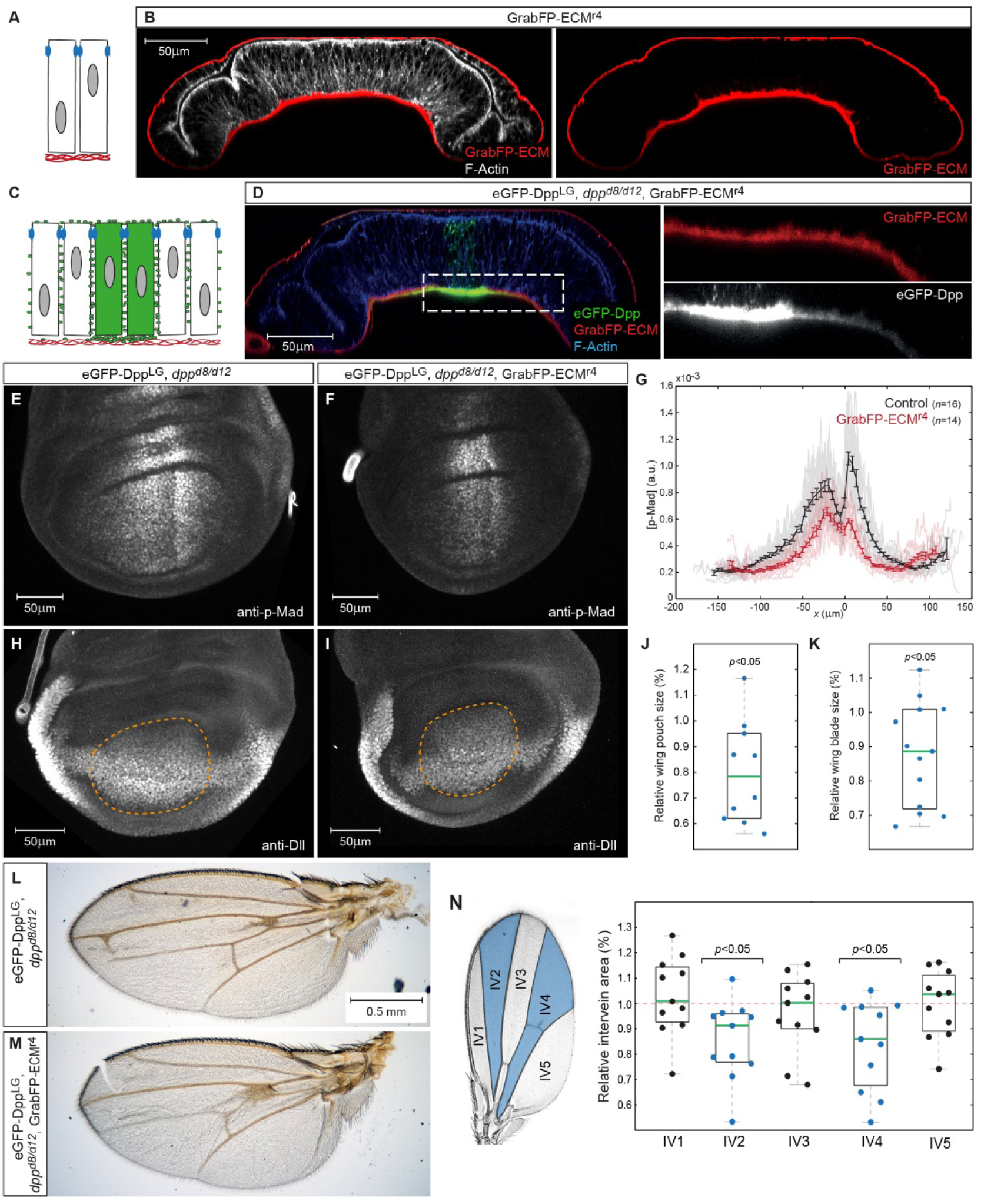
Basal Dpp is required to control wing size. **A**, Schematic representation of GrabFP-ECM localisation when expressed in the larval fat body. **B**, Wing disc optical cross-section of an animal expressing GrabFP-ECM in the fat body, stained for mCherry (GrabFP-ECM, red) and F-Actin (Phalloidin, white). **C**, Schematic of eGFP-Dpp immobilization in the ECM by GrabFP-ECM. **D**, Optical cross-section of a *dpp^d8/d12^* mutant wing disc rescued by eGFP-Dpp (green) and GrabFP-ECM (red) localizing to the basal lamina. Tissue outlines are visualized by F-Actin staining (blue). Magnification to the right shows strong eGFP-Dpp accumulation below Dpp source cells. **E-F**, 98-100h AEL old wing discs of the indicated genotype stained for p-Mad. **G**, Average p-Mad gradient at 98-100h AEL. **H-I**, 98-100h AEL old wing discs of the above indicated genotypes stained for Dll. The wing pouch is outlined by a dotted yellow line and quantified in (J). **J**, Relative wing pouch area of *dpp^d8/d12^* mutant wing disc rescued by eGFP-Dpp and GrabFP-ECM localizing to the basal lamina (*n*=11). K, Relative wing blade area of *dpp^d8/d12^* mutant wing disc rescued by eGFP-Dpp and GrabFP-ECM localizing to the basal lamina (*n*=14). **L-M**, Representative female wings of the genotypes indicated. **N**, Quantification of intervein area in GrabFP-ECM flies relative to control wings (*n*=11).

When GrabFP-ECM was expressed in the fat body of *dpp^d8/d12^* mutant larvae rescued with eGFP-Dpp (GrabFP-ECM_Rescue_ flies), high levels of eGFP-Dpp were immobilized in the BL underlying the Dpp source and low, graded levels were immobilized in the BL further away from the source stripe (Figure 8C-D). Hence, GrabFP-ECM can specifically trap Dpp and affect its dispersal in the BL, while Dpp dispersal in the lateral plane of the disc epithelium remains unaffected (Figure 8D).

To study the function of Dpp dispersal in the BL, we compared p-Mad signalling profiles in control discs and in discs of GrabFP-ECM_Rescue_ flies (Figure 8E-G). Wing discs of GrabFP-ECM_Rescue_ flies showed a clear reduction in p-Mad signalling range and peak levels (Figure 8E-G). The reduction in p-Mad range was accompanied by a significant reduction in wing disc pouch size (Figure 8H-J) and adult wing blade area (Figure 8K-M). These findings suggest that basally secreted Dpp and/or Dpp spreading in the BL contribute to proper Dpp signalling range and size control. However, despite a clear reduction in size, the overall patterning of the wing seemed unaffected in the GrabFP-ECM condition (Figure 8L-M) suggesting that basal Dpp is not strictly required for patterning the fly wing. Yet, quantification of the intervein areas showed that the medial region adjacent to the Dpp source is most susceptible to a reduction of Dpp signalling levels (Figure 8N).

## Discussion

Many proteins localize to specific membrane domains or organelles within a cell or a tissue, and it has been shown in several cases that proper protein localization plays a vital role in cell homeostasis (Wodarz and Nathke 2007, Mellman and Nelson 2008). However, the functional implication and the necessity of proper localization, as well as the consequences of distinct mislocalization of a given protein, are less well understood. Here, we have developed and used a novel, nanobody-based toolset, the GrabFP system, to interfere with the localization of GFP-tagged proteins along the apical-basal axis in the larval wing imaginal disc.

### The GrabFP system can interfere with protein localization

Recently, it was reported that tethering of nanobodies to specific cellular compartments can result in protein relocalization (Berry, Olafsson et al. 2016). In line with these observations, expression of the GrabFP constructs altered the subcellular localization along the apical-basal axis of the 15 different GFP-tagged cytosolic or transmembrane proteins we tested. All the different components of the GrabFP system induced drastic mislocalization of target proteins, causing the gain of a novel subcellular fraction, which was minor or absent in wild-type conditions. In addition, the GrabFP system significantly depleted the physiological subcellular fractions of two-thirds of the tested target proteins.

An interesting target that was effectively mislocalized is the transmembrane receptor Notch (Notch-YFP). Notch signalling is required for cell-cell communication and differentiation during development (Guruharsha, Kankel et al. 2012). The apical localization of Notch is conserved in different tissues and organisms, suggesting that it is crucial for Notch function (Fehon, Johansen et al. 1991, Ohata, Aoki et al. 2011, Hatakeyama, Wakamatsu et al. 2014). In particular, Notch apical localization might be necessary to allow interaction with its ligand Delta, which also localizes to the apical cell surface (Sasaki, Sasamura et al. 2007). In future studies, the GrabFP system will help to better understand the requirements for polarized distribution of signalling pathway components in different developmental contexts. In line with observations made by *Berry et al*. (Berry, Olafsson et al. 2016), the GrabFP components were in some cases themselves mislocalized due to interaction with target proteins. This was particularly relevant for GrabFP-A_Ext_ and GrabFP-B_Int_ (Figure 2-Figure Supplement 1G and Figure 3-Figure Supplement 1H), which were, presumably as a consequence, less efficient in causing target protein mislocalization.

In conclusion, the GrabFP system provides a general and convenient framework to specifically mislocalize GFP-tagged proteins, and large collections of GFP-tagged protein are available in *D. melanogaster* (Lowe, Rees et al. 2014, Lye, Naylor et al. 2014, Nagarkar-Jaiswal, Lee et al. 2015, Sarov, Barz et al. 2016). Moreover, the GrabFP system can be induced in a tissue-specific and temporally-controlled manner and thus represents a versatile tool to study the effect of forced protein mislocalization and protein function in specific subcellular compartments *in vivo*.

### Localized nanobodies to study the functional role of protein localization

Using Sqh-GFP, we have provided a first example of GrabFP implementation for functional studies on protein localization. We have initially described a role of Sqh during dorsal closure in the *Drosophila* embryo using the deGradFP system (Caussinus, Kanca et al. 2012). Tissue-specific degradation of Sqh (which leads to a failure to contract actomyosin networks) combined with laser ablation studies have now shown that amnioserosa cell constriction but not actin cable tension in the adjacent dorsal ectodermal cells autonomously drives dorsal closure (Pasakarnis, Frei et al. 2016). Similarly, the role of Sqh localization and the effect of Sqh mislocalization on epithelial cell shape can now be studied in more detail using the GrabFP toolset combined with other approaches such as laser ablation and force measurement.

### A basolateral Dpp pool is essential for patterning the *Drosophila* wing imaginal disc

We have previously used morphotrap to show that spreading of eGFP-Dpp is required for wing pouch patterning and for medial growth, while it is dispensable for lateral wing disc growth (Harmansa, Hamaratoglu et al. 2015). Based on this finding, we have used the GrabFP system to further dissect the functional role of eGFP-Dpp spreading with regard to the apical-basal axis in *Drosophila* wing disc development. We find that the vast majority of the eGFP-Dpp pool can be immobilized on the basolateral side of disc cells, indicating that Dpp spreads in the basolateral intercellular space. In line with this, functional interference with Dpp spreading in the basolateral compartment only (GrabFP-B_Ext_) suggests that the patterning function of the Dpp gradient is brought about to a large extend by Dpp spreading in the lateral plane of the wing disc epithelium. Growth control, in contrast, is influenced by Dpp dispersing in both the lateral and in the basal plane. These results are based on the findings that restricting basolateral Dpp dispersal (using GrabFP-B_Ext_) strongly impairs pattern and size while immobilizing eGFP-Dpp in the BL (using GrabFP-ECM) only impairs the size of the *Drosophila* wing.

Our finding of a prominent role of the basolateral compartment in Dpp spreading is interesting with regard to the mechanism of gradient formation and, at the same time, raises several new questions. Dpp gradient formation in the *Drosophila* wing disc remains a paradigm to study morphogen dispersal and several mechanisms for morphogen gradient formation have been suggested, operating in different extracellular environments (for a recent review see Akiyama and Gibson 2015). These proposed mechanisms include free extracellular diffusion in the wing disc lumen (Zhou, Lo et al. 2012), restricted extracellular diffusion in the lateral plane of the epithelium (Belenkaya, Han et al. 2004, Akiyama, Kamimura et al. 2008, Schwank, Dalessi et al. 2011), and active transport by actin-based filopodial extensions called cytonemes along the apical surface of DP cells (Hsiung, Ramirez-Weber et al. 2005). While the formation of the functional Dpp gradient in the lateral compartment is compatible with a restricted extracellular diffusion mechanism, it is not as easily compatible with the formation of a functional Dpp gradient via free diffusion in the lumen or with a key role of apical cytonemes in Dpp readout. Since we have not been able to visualize apical cytonemes, neither in wild type discs nor in disc, in which Dpp spreading along the basolateral side was blocked, we cannot make firm statements about a direct involvement of apical cytonemes in either situation.

In line with Dpp gradient formation via restricted extracellular diffusion, several studies highlighted that Dpp morphogen receptors (Lecuit, Brook et al. 1996, Lecuit and Cohen 1998, Lander, Nie et al. 2002, Crickmore and Mann 2006) and interaction partners (e.g. Dally, Belenkaya, Han et al. 2004, Akiyama, Kamimura et al. 2008) found along the extracellular surface of target cells crucially influence morphogen gradient shape. Therefore, future studies will need to investigate the localization and the effect of forced mislocalization of Dpp receptors and interaction partners on Dpp dispersal and gradient formation.

Furthermore, using a GrabFP toolset based on nanobodies or protein binders against other fluorescent proteins (Brauchle, Hansen et al. 2014), the Dpp ligand and the Dpp receptors or interaction partners could be localized to different compartments and the effect of such altered localisation could confirm or refute emerging hypotheses. Of course, it will be of critical importance to complement the results obtained using the GrabFP system with functional studies interfering with trafficking and secretion of Dpp in producing cells.

## Material and Methods

### Fly strains

The following fly lines were used: *y^1^w^1118^* (wild type), Crb-GFP (Y. Hong, Huang, Zhou et al. 2009). *dpp-LG86Fb* (K. Basler, Yagi, Mayer et al. 2010), *LOP-eGFP-Dpp* and *LOP/UAS-morphotrap* (*Harmansa, Hamaratoglu et al. 2015*), *tub>CD2,Stop>Gal4* (F. Pignioni), *sqh^A×3^* and *sqhSqh-GFP* (R. Karess) The fly stocks Dally-YFP, Dlp-YFP, Nrv1-YFP, Nrv2-YFP, NrxIV-YFP, Arm-YFP, αCat-YFP, Hts-YFP, Notch-YFP, Ed-YFP, PMCA-YFP have been obtained from the KYOTO Stock Center (DGRC) in Kyoto Institute of Technology. The fly line Fat-GFP is described in (Sarov, Barz et al. 2016) and obtained from the VDRC stock center. *r4-Gal4* was obtained from Bloomington (BL33832). *nub-Gal4, ptc-Gal4, hh-Gal4, dpp^d8^* and *dpp^d12^* are described on FlyBase (www.flybase.org).

### Genotypes by Figure

Figure 1: **C**, *nub-Gal4 / LOP/UAS-morphotrap;* **D**, *w; nub-Gal4 / LOP/UAS-GrabFP-A_Ext_;* **E**, *w; nub-Gal4 / LOP/UAS-GrabFP-B_Ext_;*

Figure 2: **B**, *LOP/UAS-GrabFP-A_Ext_ / +; NrxIV-YFP / hh::Gal4;* **C**, *LOP/UAS-GrabFP-A_Ext_ / +; Dlp-YFP / hh::Gal4;* **D**, *LOP/UAS-GrabFP-B_Ext_ / +; Crb-GFP / hh::Gal4;* **E**, Notch-YFP / + ; *LOP/UAS-GrabFP-B_Ext_ / +; hh::Gal4 / +;* **F**, *LOP/UAS-GrabFP-B_Ext_ / Ed-YFP; hh::Gal4 /* +

Figure 3: **B**, *Fat-GFP / ptc::Gal4 ; LOP/UAS-GrabFP-B_Ext_ /* +; **C**, *ptc::Gal4 / + ; LOP/UAS-GrabFP-B_Ext_* / αCat-YFP; **D**, *Nrv1-YFP / ptc::Gal4 ; LOP/UAS-GrabFP-A_Ext_* / +; **E**, *Nrv2-YFP / ptc::Gal4 ; LOP/UAS-GrabFP-A_Ext_* / +; **F**, *Hts-YFP / ptc::Gal4 ; LOP/UAS-GrabFP-A_Ext_* / +;

Figure 4: **A**, *sqh^A×3^ / + ; sqhSqh-GFP*; **B**, *ptc::Gal4 / + ; LOP/UAS-GrabFP-B_Int_* / +; **D**, *sqh^A×3^ / + ; sqhSqh-GFP / ptc::Gal4 ; LOP/UAS-GrabFP-B_Int_* / +; **E-F**, *sqh^A×3^ / Y ; sqhSqh-GFP / ptc::Gal4 ; LOP/UAS-GrabFP-B_Int_* / +

Figure 5: **A-B**: *w; LOP-eGFP-Dpp / +; dpp-LG86Fb* / +; **C-E**: yw hsFlp; *tub>CD2,Stop>Gal4, LOP-eGFP-Dpp / UAS::morphotrap; dpp-LG86Fb* / +

Figure 6: **A**: *w; LOP-eGFP-Dpp / +; dpp-LG86Fb* / +; **B**: *w; LOP-eGFP-Dpp / UAS::morphotrap; dpp-LG86Fb / hh-Gal4*; **C**: *w; LOP-eGFP-Dpp / UAS::GrabFP-B_Ext_; dpp-LG86Fb / hh-Gal4*; **D**: *w; LOP-eGFP-Dpp / UAS::GrabFP-A_Ext_; dpp-LG86Fb / hh-Gal4*;

Figure 7: **A,E,I,M**: *w; LOP-eGFP-Dpp, dpp^d12^ / dpp^d8^; dpp-LG86Fb* / +; **B,J,N**: *w; LOP-eGFP-Dpp, dpp^d12^ / UAS::GrabFP*-AB_Ext_, dpp^d8^; *dpp-LG86Fb / hh-Gal4*; **F,K,O**: *w; LOP-eGFP-Dpp, dpp^d12^ / UAS::GrabFP*-B_Ext_, dpp^d8^; *dpp-LG86Fb / hh-Gal4*;

Figure 8: **B**: *w; UAS-GrabFP-ECM / +; r4-Gal4* / +; **D,F,I,M**: *w; LOP-eGFP-Dpp, dpp^d12^ / UAS-GrabFP-ECM, dpp^d8^; r4-Gal4 / dpp-LG86Fb*; **E,H,L**: *w; LOP-eGFP-Dpp, dpp^d12^ / dpp^d8^; dpp-LG86Fb*/ +

### Molecular cloning

The following constructs were created using standard molecular cloning techniques.

**GrabFP-B_Ext_ - *pUASTLOTattB_VHH-GFP4::Nrv1::TagBFP***. The TagBFP (Evrogen) coding sequence was inserted between the first and the second exon of the *nervana 1* (Nrv1, FlyBase ID: FBgn0015776) cDNA (BDGP DGC clone LD02379). The vhhGFP4 coding fragment (Saerens, Pellis et al. 2005) was inserted at the C-terminal end of Nrv1::TagBFP. A *Drosophila* Kozak sequence (CAAA) was added and subsequently vhhGFP4::Nrv1::TagBFP was inserted into the multiple cloning site (MCS) of the pUASTLOTattB vector (Kanca, Caussinus et al. 2014).

**GrabFP-B_Int_ - *pUASTLOTattB_TagBFP::Nrv1::vhhGFP4***. To generate a basolateral GrabFP construct that exposes the nanobody to the cytosol, tagBFP was exchanged with the vhhGFP4 coding sequence.

**GrabFP-A_Ext_ - *pUASTLOTattB_VHH-GFP4::T48-Baz::mCherry***. The HA-tag was replaced by vhhGFP4 in the T48-HA plasmid (obtained from M. Leptin, Kölsch, Seher et al. 2007). mCherry was inserted at the C-terminal end of vhhGFP4::T48. In addition, the 2316 base pair minimal apical localization sequence of Bazooka (obtained from A. Wodarz, Krahn, Klopfenstein et al. 2010) was attached C-terminally to mCherry. A Drosophila Kozak sequence (CAAA) was added when inserting vhhGFP4::T48-Baz::mCherry into the MCS of the pUASTLOTattB vector (Kanca, Caussinus et al. 2014).

**GrabFP-A_Int_ - *pUASTLOTattB_mCherry::T48-Baz::vhhGFP4***. To switch the topology we exchanged the mCherry with the vhhGFP4 coding region, resulting in orientation of the nanobody into the cytosol.

**GrabFP-ECM - *pUASTLOTattB_VHH-GFP4::Vkg::mCherry***. vhhGFP4 and mCherry coding sequences, separated by a short linker region, were inserted between the 1^st^ and 2^nd^ exon in the Vkg full-length plasmid (obtained from L. Ashe, Wang, Harris et al. 2008). This insertion site was chosen, since a viable Vkg GFP-trap line exists which carry an exogenous GFP exon at this position (Morin, Daneman et al. 2001). Finally the vhhGFP4::Vkg::mCherry construct was inserted into the MCS of the pUASTLOTattB vector (Kanca, Caussinus et al. 2014).

All transgenes were inserted by phiC31-integrase-mediated recombination into the 35B region on the 2nd chromosome and the 86Fb region on the 3^rd^ chromosome. The obtained transgenic flies respond to both, LexA and Gal4 transcriptional activators. By crossing with Cre^y^ expressing flies one of the response elements can be removed in a mutually exclusive manner. The excision was screened for by PCR as described in Kanca, Caussinus et al. 2014.

### Staging of larvae and dataset creation

Generally, third instar wandering larvae were dissected and used for analysis. However, for quantification of expression profiles (Figure 7A-H and Figure 8E-G)) or pouch size (Figure 7I-L and Figure 8H-J) larvae were staged to 98-100 hours after egg laying (AEL) as described before (Hamaratoglu, de Lachapelle et al. 2011, Harmansa, Hamaratoglu et al. 2015). Only male larvae were included in this analysis, positively selected by the presence of the transparent genitalia disc.

### Immunostaining and image acquisition

Immunostaining of larval wing imaginal discs was performed as described previously (Harmansa, Hamaratoglu et al. 2015). Staged quantitative data sets were always processed together using identical solutions and mounted on the same microscopy slide (using brains as spacers) to reduce variation. All data stacks (with slices every 1μm) of a quantitative data set were acquired within the same microscope session under imaging conditions within the linear range of the fluorescent signal obtained.

For high resolution imaging along the z-axis (optical-cross sections of wing discs) discs were mounted using double sided tape as spacer to avoid squeezing of the discs and to preserve their morphology. To obtain maximum resolution along the z-axis stacks were acquired with sections every 0.17μm.

### Antibodies

rabbit (rb)-anti-mCherry (1:5000, gift from E. Nigg), rb-anti-tRFP (1:2000, Evrogen, #AB233), mouse (m)-anti-Dlg (4F3, 1:500, DSHB, University of Iowa), rb-anti-phospho-Smad1/5 (1:300; Cell Signaling, 9516S), guinea pig (gp)-anti Dll (1:2000, a gift from R. Mann), m-anti-Wg (4D4-s; 1:120; DSHB, University of Iowa); m-anti-Ptc (Apa1-s; 1:40; DSHB, University of Iowa). Secondary antibodies from the AlexaFluor series were used at 1:750 dilution with the exception of Alexa405-anti-rb which was used at 1:500 dilution. CF405S-anti-gp was used at 1:1000 dilution (Sigma Aldrich).

### Image processing

Image data was processed and quantified using ImageJ software (National Institute of Health). Optical cross-sections were computed using the section tool in Imaris software (Bitplane).

For improved resolution, datasets in Figures 1-4 were deconvolved using the Huygens Remote Manager software (Ponti et al., 2007). Quantification of protein localization along the A-B axis is explained in the next section. Average expression profiles were obtained using the WingJ software (Schaffter 2014) (http://tschaffter.ch/projects/wingj/) as done previously (Harmansa, Hamaratoglu et al. 2015). In brief, we made use of Wg and Ptc stainings marking the D/V and the A/P boundary, respectively. Profiles were then extracted up to the edge of the wing disc with a 30% offset in the dorsal compartment along a line parallel to the D/V boundary (see (Hamaratoglu, de Lachapelle et al. 2011, Harmansa, Hamaratoglu et al. 2015)). Plotting of the average profiles was done in Matlab software (Mathworks) using the WingJ Matlab toolbox.

### Extraction of concentration profiles along the apical-basal axis

In order to quantify relative protein levels in the apical versus the basolateral compartment we acquired high z-resolution stacks (as described in the imaging section) of multiple wing discs stained for the junctional marked discs-large (Dlg). From these discs we obtained optical cross-sections in the dorsal compartment parallel to the D/V boundary using the “reslice” option in Fiji software (ImageJ, National Institute of Health) (see Figure 1-Figure Supplement 3A). From these cross-sections we extracted the fluorescent intensity profiles of Dlg and the protein of interest in a rectangular region of 114×16μm using the “plot profile” function in ImageJ (see Figure 1-Figure Supplement 3B). We used the junctional peak of the Dlg profile to align the individual target profiles of different discs. To correct for variation between profiles from different discs we (1) subtracted the background fluorescence observed in the disc lumen (minimal fluorescence intensity observed in the luminal region) and (2) subsequently normalized individual profiles to one. Average profiles were calculated in Excel software (Microsoft) and plotted in Matlab software (Matworks). In the depicted plots, we only included signal from the DP region and excluded signal from the PPM (see Figure 1-Figure Supplement 3C-D). The peak of the average Dlg profile plus and minus 1.0μm (marked by a blue bar) was defined as the junctional plane and the border between the apical and the basolateral compartment. Error bars show the standard error.

In order to quantify the absolute fluorescence levels of the proteins of interest in the apical versus the junctional and the basolateral compartments (as in Figure 2G-H and Figure 3G-H) we plotted the average fluorescence values observed in the different compartments after background subtraction.

### Statistics and data representation

Sample size was chosen large enough (n≥5) to allow assessing statistical significance using a two-sided Student’s *t*-test with unequal variance (* *p*≤0.05, ** *p*≤0.005, *** *p*≤0.0005). Sample number and *p*-values are indicated in either the figure or the figure legend for each experiment. n-numbers indicate biological replicates, meaning the number of biological specimens evaluated (e.g. the number of wing discs or wings). In boxplot graphs outliers are indicated by a red cross (e.g. Figure 1-Figure Supplement 2) and were excluded from further computation.

In the A-B concentration profiles (e.g. Figure 2B-F) bold lines represent average fluorescent values and error bars correspond to the standard error. Bold lines in the P-Mad expression profiles (Figure 7D,H and Figure 8G) indicate the arithmetic mean and the error bars show the standard deviation; individual profiles used for the analysis are shown light-coloured. In box plots individual data points are shown and the centre value represents the media while the whiskers mark the maximum and minimum data points.

## Acknowledgments

We thank F. Hamaratoglu, K. Basler and G. Pyrowolakis for discussions and input on the project, the Biozentrum Imaging Core Facility for mA_Int_enance of the microscopes and support. We are grateful to M. Leptin, A. Wodarz, L. Ashe, E. Nigg, Y. Hong and R. Mann for flies and reagents.

## Author Information

The authors declare no competing financial interests. Correspondence and requests for materials should be addressed to M.A.

## Figure Legends

**Figure 1-Figure Supplement 1.**
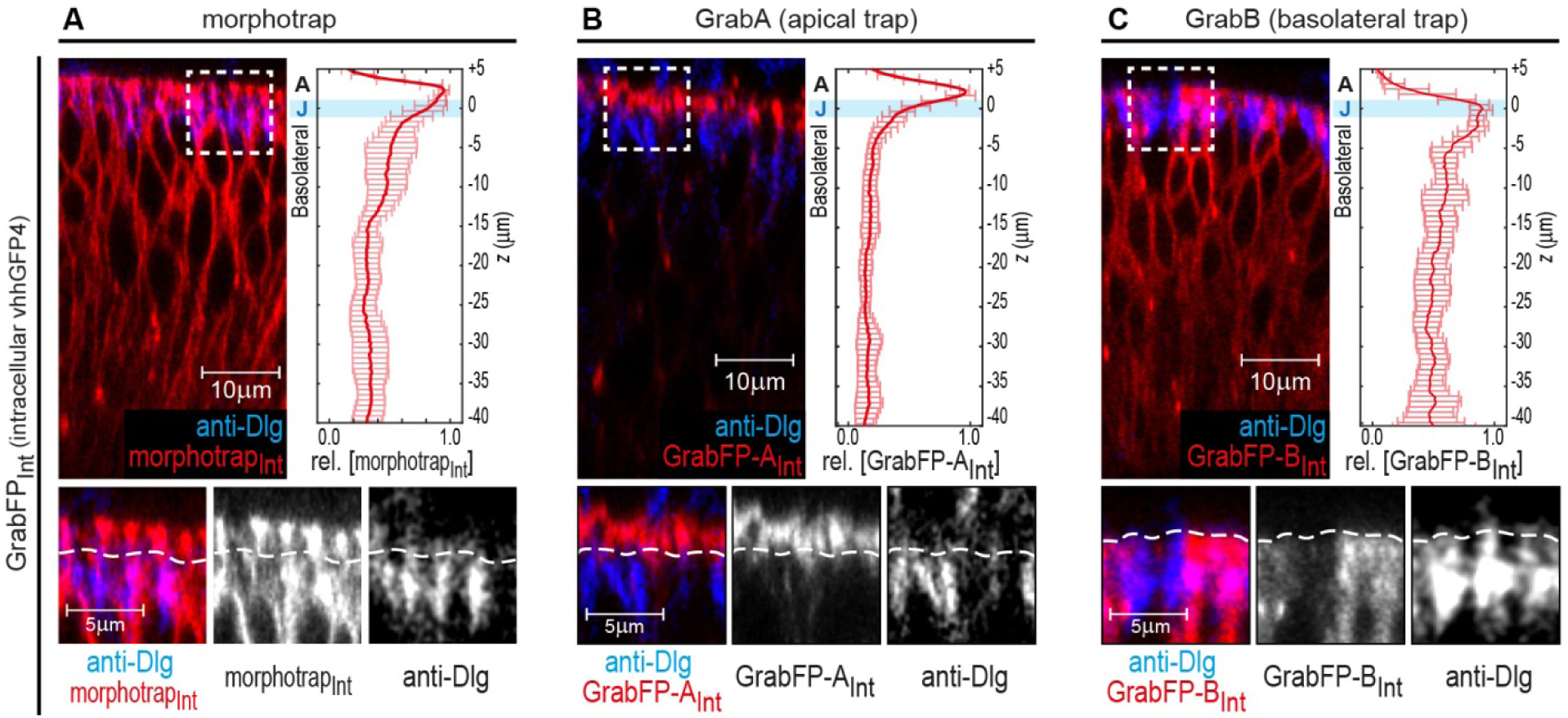
Localization of the GrabFP_Intra_ tools. **A-C**, Optical cross-sections of wing discs expressing the intracellular versions of morphotrap_Int_ (A), GrabFP-A_Int_ (B) and GrabFP-B_Int_ (C) in the disc proper (*ptc::Gal4*). The GrabFP constructs are shown in red and the junctions are visualized by staining for Dlg (blue). The junctional level is marked by dotted lines in the magnifications (bottom). Quantification of protein localization along the A-B axis is shown in the graphs to the right. (morphotrap_Int_ *n*=6, GrabFP-A_Int_ *n*=7, GrabFP-B_Int_ *n*=6, error bars represent the standard deviation). For details on the quantification see method section and Figure 1-Figure Supplement 3.

**Figure 1-Figure Supplement 2.**
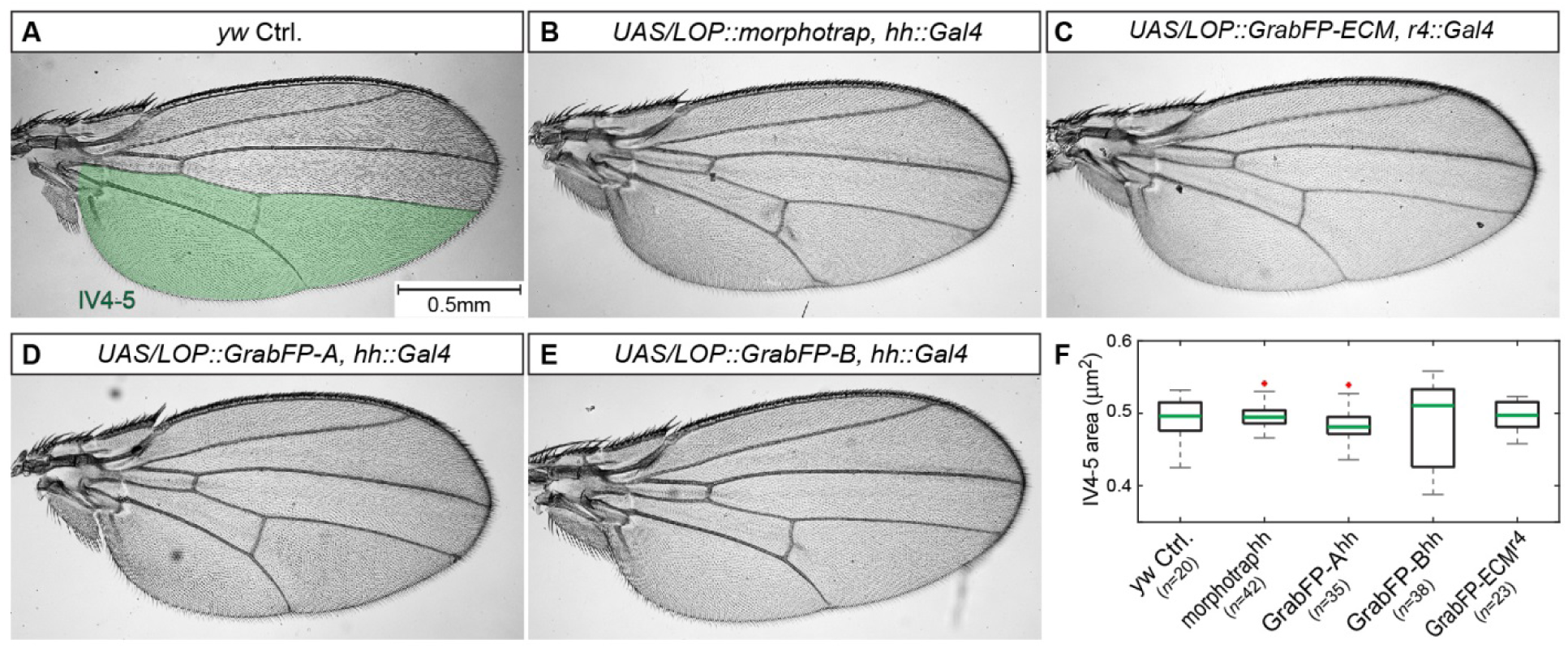
Expression of the GrabFP system allows normal wing development. **A-E**, Male wings of indicated genotypes. Expression of the GrabFP_Ext_ tools (using *hh::Gal4* (A-B, D-E) or *r4::Gal4* (C)) does not interfere with wing development and yields viable and fertile flies. Solely expression of GrabFP-A_Ext_ in the posterior compartment results in slightly rounder wing shape (compare A to D). **F**, Quantification of intervein area between vein 4 and the posterior wing margin (IV4-5), as marked in (A). None of the genotypes showed significantly reduced or increased wing blade area due to the expression of the GrabFP tools. Significance was assessed using a two-sided Student’s *t*-test with unequal variance (*p*-values: morphotrap *p*=0.59, GrabFP-A^hh^ *p*=0.26, GrabFP-B^hh^ *p*=0.61, GrabFP-ECM^r4^ *p*=0.55, outliers are marked by a red cross).

**Figure 1-Figure Supplement 3.**
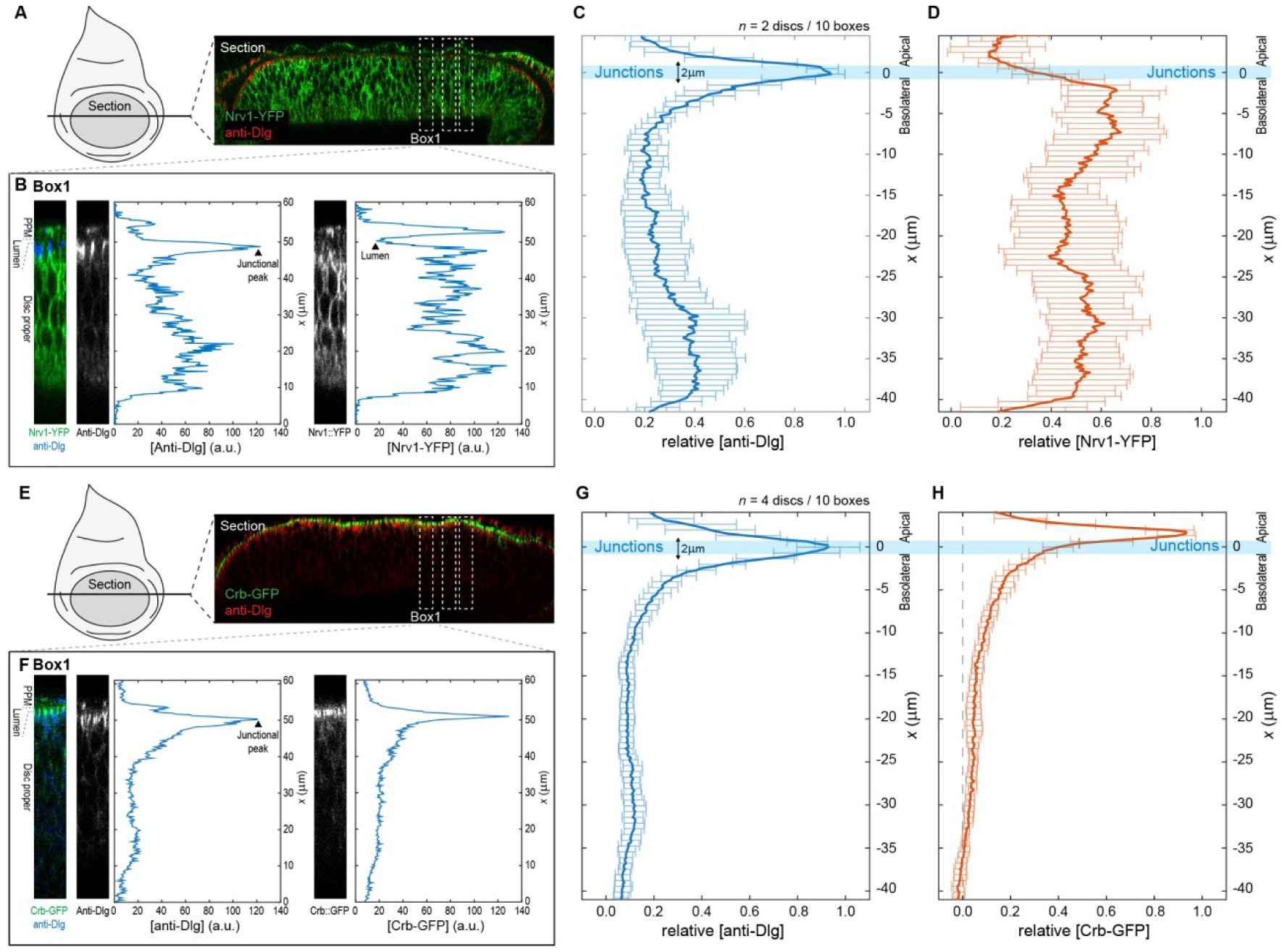
Quantification and analysis of protein distribution along the A-B axis. Procedure for obtaining relative concentration profiles along the A-B axis of DP cells for the basolateral marked Nrv1-YFP (A-D) and the apical marked Crb-GFP (E-H). **A**, Optical cross-section of a wing disc expressing Nrv1-YFP (green) and stained for Dlg (blue) as obtained when using the reslice function in ImageJ (NIH). **B**, Single fluorescence intensity profiles of anti-Dlg and Nrv1-YFP fluorescence extracted from a rectangular area of 16μm width (e.g. Box1) using the plot profile function of ImageJ. **C-D**, Individual profiles as extracted in (B) were aligned according to the position of the junctional peak of the Dlg signal and merged to average concentration profiles. Average profiles of *n*=2 discs/10 sections are shown for Dlg (C) and Nrv1-YFP (D). The junctions are defined as the region 1μm above and below the average Dlg peak (light blue bar). Nrv1 localization is restricted to the basolateral compartment, and indeed our quantifications show that Nrv1-YFP levels are high along the basolateral compartment but drop at the junctions. **E-H**, Similar steps as in (A-D) for the extraction of average Crb-GFP profiles. Crb is a determinant of apical compartment identity and exclusively localizes to the apical compartment as visualized by the quantification.

**Figure 2-Figure Supplement 1.**
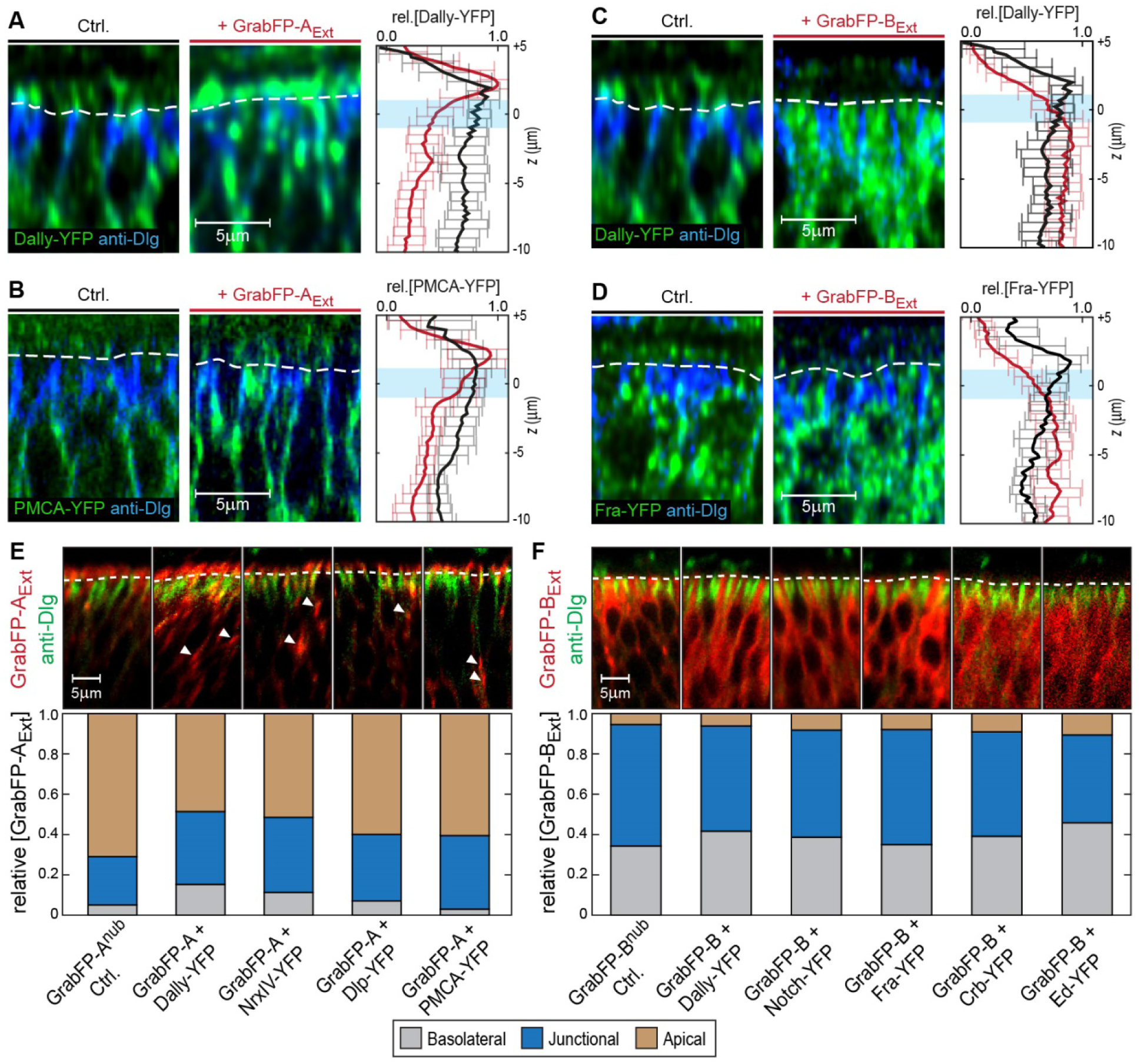
Examples of target protein mislocalization using the GrabFP_Extra_ system. **A-B**, Effects of GrabFP-A_Ext_ expression on the localization of Dally-YFP (A) and PMCA-YFP (C). Optical cross-sections of target proteins alone (Ctrl., left) or co-expressed with GrabFP-A_Ext_ (middle). Quantification of target-protein levels (left) in the absence (black) or the presence of GrabFP-A_Ext_ (red). C-D, Localization of Dally-YFP (C) and Fra-YFP (D) in control conditions (left) and when co-expressed with GrabFP-B_Ext_ (middle), quantification is shown to the left. **E-F**, Representative cross-sections (top) and quantification (bottom) of relative GrabFP-A_Ext_ (E) and GrabFP-B_Ext_ (F) localization when expressed alone (Ctrl.) or when co-expressed with YFP/GFP-tagged target protein. Target proteins of basolateral localization tend to mislocalize GrabFP-A_Ext_ (E) towards the junctional/basolateral compartment (see reduction in relative apical localization (brown)). In contrast, GrabFP-B_Ext_ (F) is more resistant to mislocalization by apically localizing target proteins and shows only slight increases in apical localization. (Sample numbers for shown quantifications in A-F: GrabFP-A_Ext_ *n*=4, +Dally *n*=10, +NrxIV *n*=6, +Dlp *n*=8, +PMCA *n*=5, GrabFP-B_Ext_ *n*=4, +Dally *n*=10, +Notch *n*=8, +Fra *n*=9, +Crb *n*=9, +Ed *n*=6)

**Figure 3-Figure Supplement 1.**
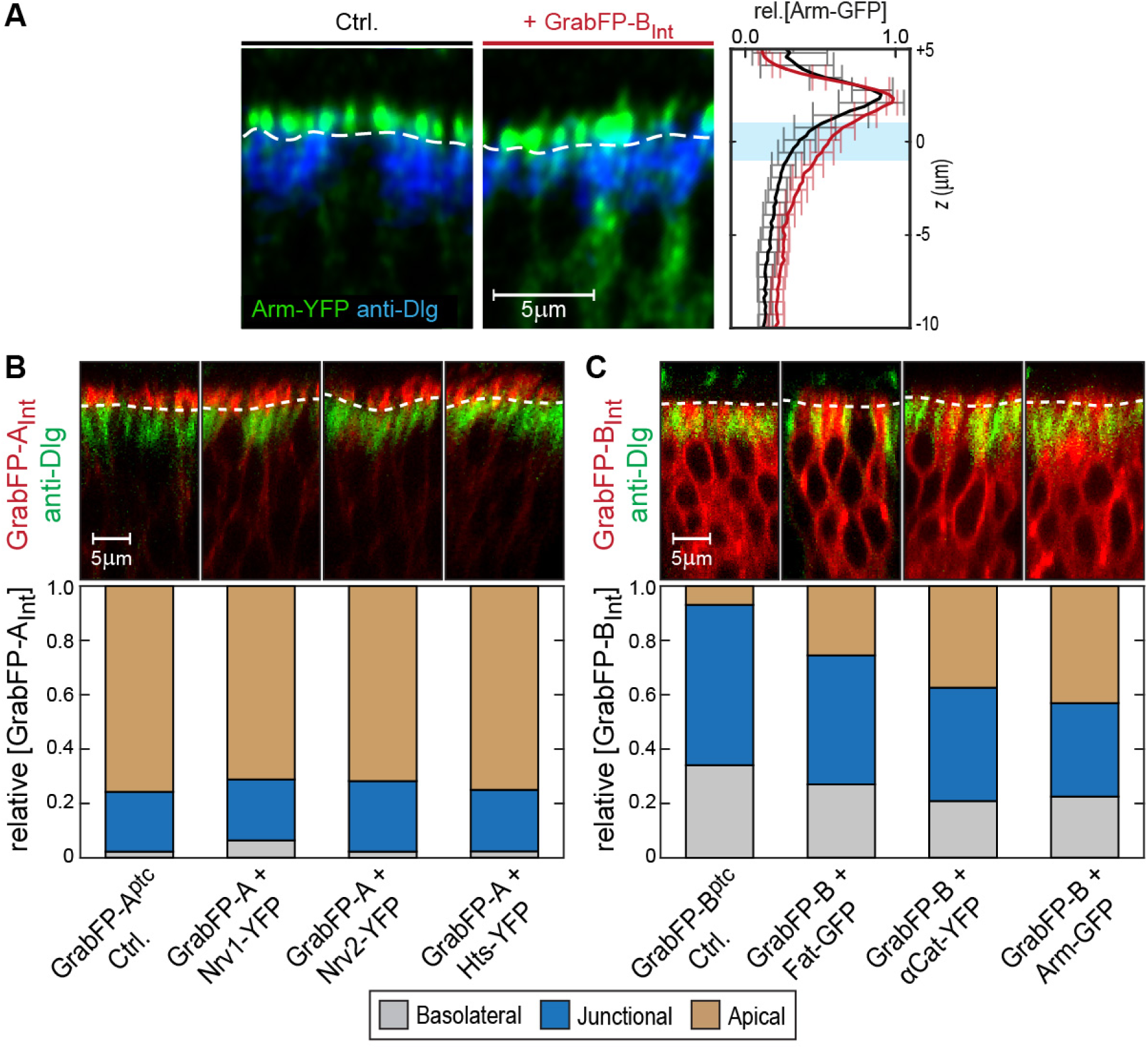
Examples of GFP/YFP-target protein mislocalization using the GrabFP_Intra_ system. Optical cross-sections of DP cells expressing Arm-YFP (Ctrl., left) and Arm-YFP together with GrabFP-B_Int_ (middle). Average Arm-YFP protein distribution along the A-B axis in the absence (black) and in the presence of GrabFP-B_Int_ (red) is plotted to the right. **B-C**, Representative cross-sections (top) and quantification (bottom) of relative GrabFP-A_Int_ (B) and GrabFP-B_Int_ (C) localization when expressed alone (Ctrl.) or when co-expressed with YFP/GFP-tagged target protein. While GrabFP-A_Int_ is robust to mislocalization by target proteins, GrabFP-B_Int_ tends to be mislocalized to the apical compartment in all three conditions tested. (GrabFP-A_Int_ Ctrl. *n*=7, Nrv1 *n*=10, Nrv2 *n*=10, Hts *n*=10, GrabFP-B_Int_ Ctrl. *n*=6, Fat *n*=10, αCat *n*=9, Arm *n*=8)

**Figure 6-Figure Supplement 1.**
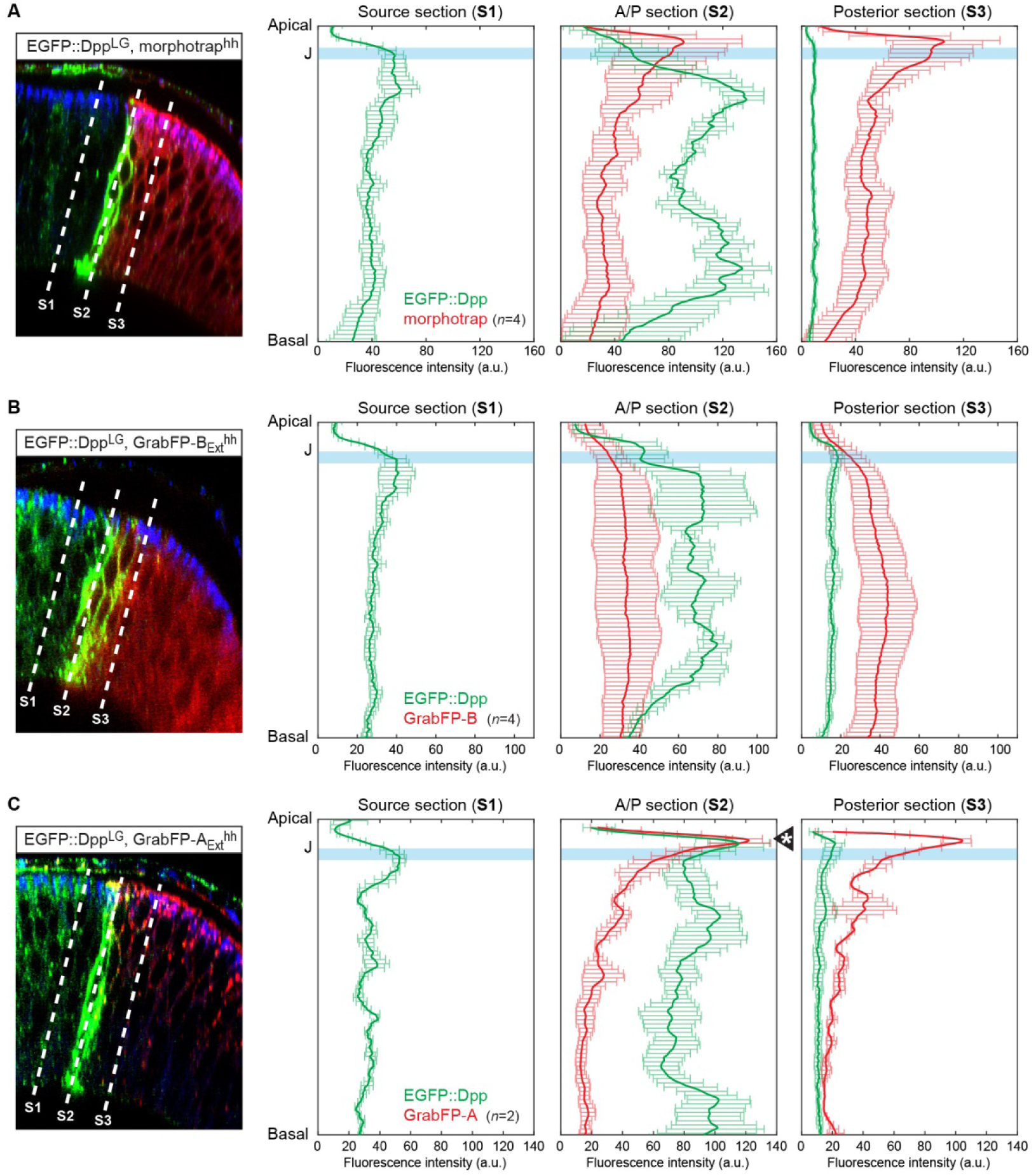
Quantification of differential eGFP-Dpp accumulation by morphotrap, GrabFP-B_Ext_ and GrabFP-A_Ext_. eGFP-Dpp immobilization pattern along the A-B axis in the posterior compartment when either morphotrap (**A**), GrabFP-B_Ext_ (**B**) or GrabFP-A_Ext_ (**C**) are expressed in posterior cells (*hh::Gal4*). Left column: Optical cross-sections as shown in Figure 5. Positions at which eGFP-Dpp and GrabFP_Ext_ localization was measured are indicated by dotted lines (S1-S3). Middle-right column: Plots of average eGFP-Dpp (green) and GrabFP_Ext_ (red) levels along the A-B axis as positions indicated in the left column. eGFP-Dpp levels are strongly increased along the A/P boundary (S2 section) in all conditions. Importantly, neither morphotrap (A, middle-right) nor GrabFP-B_Ext_ (B, middle-right) immobilize eGFP-Dpp in the apical compartment above the junctions (thick blue line). In contrast, GrabFP-A_Ext_ shows strong eGFP-Dpp immobilization in the apical compartment (arrowhead in C, middle-right). This might be due to GrabFP-A_Ext_ mediated mislocalization of basolateral eGFP-Dpp to the apical compartment. In all conditions eGFP-Dpp levels drop after the first 2-3 cell rows (green in S3) suggesting that indeed posterior GrabFP_Ext_ expression results in impaired posterior Dpp dispersal. In the plots, thick lines represent average fluorescence values and error bars show the standard deviation.

**Figure 7-Figure Supplement 1.**
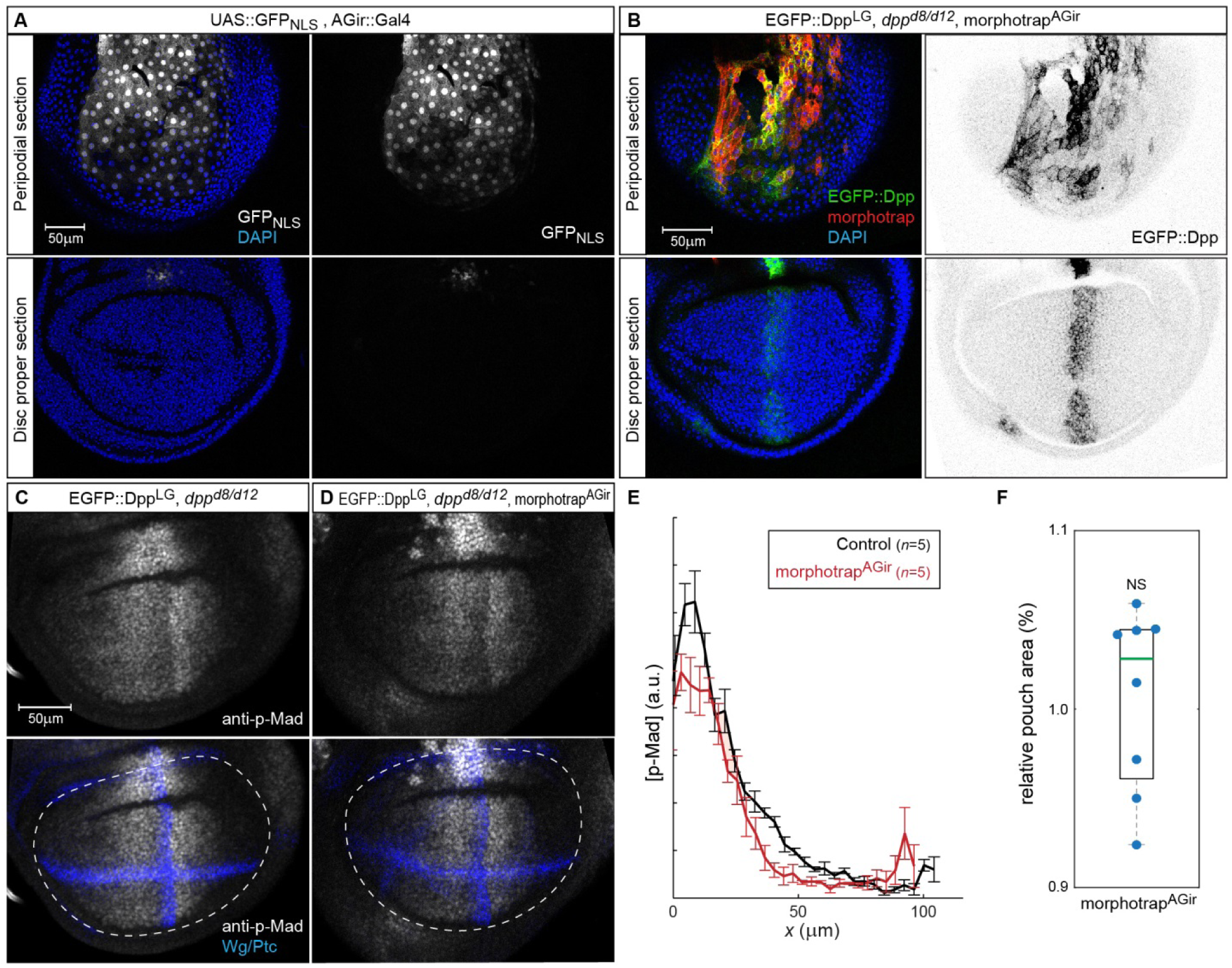
GrabAB_Ext_ expression in PPM cells interferes with luminal Dpp spreading. **A**, Expression of *UAS-GFP_NLS_* under control of the *AGir::Gal4* driver line in a third instar wing imaginal disc. The activity of *AGir::Gal4* is restricted to PPM cells and a small group of cells in the dorsal hinge region of the DP. **B**, *dpp^d8/d12^* mutant wing disc rescued by eGFP-Dpp (green) expressing morphotrap (red) in the PPM (*AGir::Gal4*). A projection of the PPM plain shows that high amounts of luminal eGFP-Dpp are immobilized along PPM cells expressing morphotrap (top), while eGFP-Dpp dispersal is undisturbed in DP cells (bottom). **C-F**, Patterning and growth are largely unaffected by morphotrap mediated immobilization of eGFP-Dpp along PPM cells. Peak levels of Dpp signalling activity, visualized by p-Mad (grey), are slightly reduced in the morphotrap condition (D) compared to controls (C, 98-100h AEL). Quantification is shown in (E). Pouch size (white dotted line in C-D), as defined by the inner Wg ring (blue in C-D), is also not affected by modifying luminal Dpp dispersal (F).

